# The *Staphylococcus aureus* serine protease-like protein B is a potent allergen in a murine asthma model

**DOI:** 10.1101/2025.09.10.675349

**Authors:** Jessica von Fournier, Marques L. Schilling, Christopher Saade, Hannes Wolfgramm, Shenja Buchholz, Susanne Neumeister, Yves Laumonnier, Henry J. McSorley, Sabrina Schulze, Leif Steil, Manuela Gesell Salazar, Matthias Sendler, Silva Holtfreter, Uwe Völker, Murthy N. Darisipudi, Barbara M. Bröker

**Author notes:** Corresponding author Jessica von Fournier, University Medicine Greifswald, Institute of Immunology, Ferdinand-Sauerbruch-Straße / DZ 7, 3rd Floor 17475 Greifswald, Germany. Conflicts of interests: The authors declare that they have no conflicts of interest.

## Abstract

**Background:** Asthma is associated with *Staphylococcus aureus* colonization. Two hypotheses were proposed to explain this phenomenon: (1) the allergic environment in asthma favors *S. aureus* colonization and (2) *S. aureus* colonization creates a pro-allergic environment. Since several *S. aureus* virulence factors, such as the serine protease–like protein (Spl) B, elicit a type 2 biased immune response, we asked whether the pathogen itself can cause asthma.

**Objective:** Test the ability of recombinant SplB of *S. aureus* to sensitize mice and induce allergic airway inflammation (AAI).

**Methods:** Mice were treated with repeated intratracheal inoculations of either catalytically active SplB or an inactive mutant. AAI was assessed by evaluating airway hypersensitivity, immune cell infiltration, cytokines, mucus production, fibrosis, and specific serum IgE. We compared the outcome between wild-type and gene-deficient C57BL/6J mice, including recombination-activating gene knockout mice (*Rag2^-/-^*), interleukin-33 knockout mice (*Il33 ^-/-^*), and protease-activated receptor 2 knockout mice (*F2rl1^-/-^*).

**Results:** Intratracheal exposure to SplB sensitized the mice and caused eosinophilic airway inflammation and hyperresponsiveness. The development of asthma required both the proteolytic activity of SplB and a functional adaptive immune system. The soluble protease sensor IL-33 was necessary for eosinophil tissue invasion, whereas the membrane-bound protease sensor PAR2 was not.

**Conclusion:** The serine protease SplB of *S. aureus* is a potent allergen. Based on this finding we propose a third mechanism to explain the relationship between *S. aureus* colonization and asthma: *S. aureus* can release allergens, such as SplB, that sensitize individuals and lead to the development of asthma.

**Key Messages:**

- The SplB protease, which is secreted by *S. aureus*, is a potent allergen that induces eosinophilic airway inflammation in mice.
- SplB’s allergenicity depends on its enzymatic activity and on the host’s adaptive immune system.
- IL-33, a soluble protease sensor, is required for eosinophilic airway inflammation but not for the SplB-specific IgE response. The membrane protease sensor PAR2 was only required for lung fibrosis.
- Bacterial proteases may be an underestimated cause of idiopathic asthma.

**Capsule summary:** The protease SplB, produced by the bacterium *Staphylococcus aureus*, induces allergic airway inflammation in mice, suggesting that *S. aureus* may promote asthma in human patients. These findings imply that bacterial allergens could play a significant role in idiopathic asthma.

## Introduction

Asthma is one of the most common chronic inflammatory lung diseases, with over 260 million patients worldwide^1^. Patients suffer from wheezing, cough, chest tightness and shortness of breath. Asthma is a heterogeneous disease, and clinicians differentiate between several asthma phenotypes, of which allergic asthma is the most common^2^. Allergic asthma is characterized by a T helper type 2 cell (Th2) immune response and symptoms are often triggered by the inhalation of airborne allergens. However, the causes of asthma development are debated^3^.

Colonization by certain pathogens, among them *Staphylococcus aureus*, is suspected to play a role in asthma development and exacerbation^4,5^. *S. aureus* is a Gram-positive opportunistic pathogen that colonizes the airways in around 30% of the adult human population^6^ Colonization with *S. aureus* is positively correlated with asthma and also associated with other atopic diseases like atopic dermatitis and chronic rhinosinusitis^7,8^. Two mutually non-exclusive scenarios have been proposed to explain the association between *S. aureus* colonization and asthma: (1) the Th2 immune response that characterizes asthma facilitates *S. aureus* colonization and (2) *S. aureus* colonization creates a Th2 microenvironment that promotes allergic sensitization towards harmless antigens^5^.

The immune response towards prominent human asthma allergens like house dust mite (HDM), pollen and cockroach is well-defined^10–12^. Many of these allergens contain proteases, which provoke a type 2 immune response by cleaving tight junction proteins and activating cell-surface protease sensors, such as the protease-activated receptor 2 (PAR2)^13–17^. Moreover, several protease allergens cleave and thereby activate interleukin (IL)-33, an alarmin that is released by epithelial cells upon cellular stress and cell death^18^.

*S. aureus* can produce a plethora of virulence factors, some of which have been linked to a type 2 response in humans, such as the serine protease-like proteins (Spls)^9^. The Spls are a group of serine proteases (commonly named SplA-SplF) that are secreted by *S. aureus* and encoded in its genome on the νSaβ genomic island^19–21^. The Spls share amino acid sequence homology between 43.9 and 94.6% and have each distinctive amino acid cleavage motifs^21,22^. In a mouse sepsis model, they act as virulence factors and increase mortality^23^. Moreover, SplD and SplF cause asthma in an acute mouse asthma model. However, the mechanisms underlying their allergenicity remain poorly defined^24^. In humans who are naturally exposed to *S. aureus*, the Spls elicit a type 2-biased immune response in T cells and B cells^25^. The antibody response directed against the Spls is skewed towards IgE and IgG4 and in asthmatics the Spl-specific IgE antibody titers are increased^26^. Interestingly, measurable anti-SplB IgE titers were reported even in many clinically healthy individuals^26^, making this protease an intriguing target for further investigation.

To shed light on the interplay between *S. aureus* and airway allergy, we have studied the *S. aureus* protease SplB. We show that SplB acts as a potent allergen in a mouse model of allergic airway inflammation (AAI). SplB alone – without adjuvant – is sufficient to cause eosinophilic inflammation. The development of eosinophilic asthma depends on the adaptive immune system and on SplB’s protease activity. IL-33 is necessary for the onset of symptoms, but PAR2 is not. Therefore, we propose a third scenario for the relationship between *S. aureus* and asthma: (3) *S. aureus* secretes allergens that can trigger asthma development and exacerbation.

## Methods

### Recombinant proteins

Recombinant SplB was produced in the *S. aureus* strain RN4220 using the expression vector pTripleTREP^27^. The sequence of SplB was derived from the *S. aureus* strain USA300 FPR3757 (Fig. E1 in the Online Repository). An amino acid exchange (S193A) was introduced into the catalytic triad of SplB to generate a SplB mutant (SplB mut) without catalytic activity. The recombinant proteins lack the N-terminal signal peptide, as this is cleaved off during secretion by the host *S. aureus* strain, and they harbor a C-terminal Twin-Strep-tag®.

### Mice

Animals were maintained in a 12-hour/12-hour light/dark cycle and had access to water and food *ad libitum*. Animal experiments were approved by the responsible authority, the Landesamt für Landwirtschaft, Lebensmittelsicherheit und Fischerei Mecklenburg-Vorpommern (LALLF MV; AZ 7221.3-1-037/20). All animal experiments were carried out in accordance with the European Union Directive 2010/63/EU, the German Animal Welfare Act (Tierschutzgesetz), and the German Animal Welfare Ordinance on the Protection of Animals Used for Experimental Purposes (TierSchVersV). The following mouse strains were used: C57BL/6J, C57BL/6J-Rag2^em3Lutzy^/J (*Rag2^-/-^*), C57BL/6J *Il33^Gt/Gt^* (*Il33^-/-^*)^28^ and B6.Cg-F2rl1tm1Mslb/J (*F2rl1^-/-^*)^29^. Mice were purchased from Janvier (C57BL/6J), JAX (C57BL/6J *Rag2^-/-^* (strain 033526); C57BL/6J *F2rl1^-/-^* (strain 004993)) or they were bred in-house (C57BL/6J *Il33^-/-^*, originally from Texas A&M Institute for Genomic Medicine (USA)). The mice were certified as *S. aureus*-free. All mice used in animal experiments were female and 8-12 weeks old when treatment started.

### Mouse model of allergic airway inflammation (AAI)

Mice were put under anesthesia (75 mg/kg ketamine + 5 mg/kg xylazine) and treated intratracheally (i.t.) with PBS, SplB (10, 25, or 50 µg), SplB mut (25 µg) or OVA (50 µg) on days 0, 7, 14 and 21. Notably, no adjuvant or intraperitoneal (i.p.) priming was used in either of the groups. Proteins were dissolved in 50 µL PBS. In some experiments, the IL-33 inhibitor HpARI2 (10 µg), the PAR2-blocking monoclonal antibody (mAb) SAM-11 (10 ng; ThermoFisher Scientific, USA; 35-2300) or a monoclonal IgG isotype antibody (10 ng; ThermoFisher Scientific, USA; 02-6200) were co-applied i.t. with SplB. On days 0, 7 and 14, approximately 100 µL blood was taken from the retroorbital plexus prior to the i.t. application. 72 hours after the fourth treatment, mice were anesthetized and the airway hyperreactivity (AHR) was measured. Afterwards the mice were sacrificed with an overdose of isoflurane.

### Flow cytometry

A comprehensive list of antibodies used for flow cytometric analyses is provided in the Methods section of the Online Repository, and representative gating strategies are shown in Figures E2-E5.

### Statistics

All datasets were tested for normality using the Shapiro–Wilk test. Ordinary one-way ANOVA followed by Tukey’s multiple comparisons test was used if all datasets were normally distributed, otherwise the Kruskal–Wallis test followed by Dunn’s multiple comparisons test was applied. AHR measurements were analyzed by calculating the area under the curve (AUC) and the AUC values were compared using the Kruskal–Wallis test followed by Dunn’s multiple comparisons test. A p value < 0.05 was considered significant. Statistical analyses were performed with GraphPad Prism v8.0.1. Data are shown as median with interquartile range if more than one group consisted of less than 4 animals, and as mean ± SD otherwise.

A more detailed description of the materials and methods used in this study can be found in the Methods section in this article’s Online Repository.

## Results

### Catalytically active SplB acts as a potent allergen in murine allergic airway inflammation (AAI)

First, we established a protocol for AAI in female C57BL/6J WT mice. For sensitization we applied SplB i.t. on days 0, 7 and 14. On day 21 the animals were challenged with a fourth i.t. dose of SplB (Fig. 1A). SplB was tested at different doses. No adjuvant was used. Even at the lowest dose (10 µg/application), the protease elicited prominent eosinophilic lung inflammation, accompanied by increased airway hyperreactivity (AHR) (Fig. E6 in the Online Repository). We decided to treat mice with 25 µg/dose except for experiments assessing the effects of an inhibitor, when a lower amount of 10 µg/dose was used.

**Figure 1.**
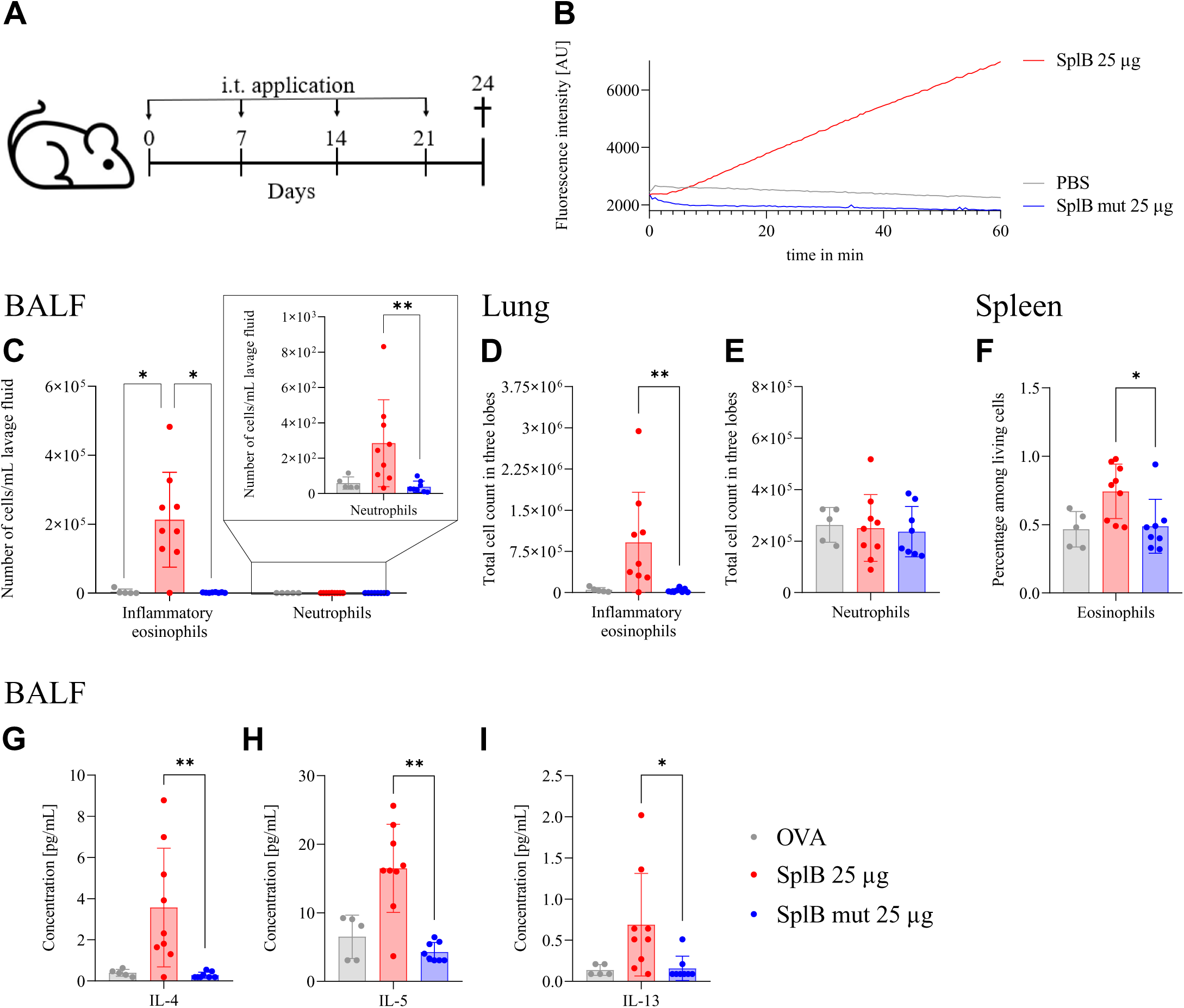
Catalytic activity of SplB is essential to cause AAI. Female WT C57BL/6J mice were treated with active SplB or the catalytically inactive SplB mut; OVA-treated mice served as control. **A**, Mouse model of AAI. Mice received weekly intratracheal (i.t.) applications of either PBS or different doses of SplB and were sacrificed 72 hours after the fourth application. **B**, SplB mut is catalytically inactive. The catalytic activity of SplB and SplB mut was determined with the synthetic peptide substrate Ac-VEID-MCA. After substrate addition, fluorescence was measured over 60 min. PBS served as negative control. One out of three similar experiments is depicted. **C**, Numbers of inflammatory eosinophils and neutrophils in the BALF measured by flow cytometry. **D** and **E**, Numbers of inflammatory eosinophils (D) and neutrophils (E) in the lung measured by flow cytometry. **F**, Percentage of eosinophils in the spleen measured by flow cytometry. **G-I**, Concentrations of IL-4 (G), IL-5 (H) and IL-13 (I) in the BALF measured by CBA. n = 5-9. Results are presented as mean ±SD. *p < .05 and **p < .01. Significance was determined using a Kruskal–Wallis test followed by Dunn’s multiple comparisons test.

A functional adaptive immune system was required for the development of AAI because *Rag2^-/-^* mice, who lack T- and B cells, did not develop any signs or symptoms of AAI (Fig. E7 in the Online Repository). Type 2 cytokine levels in SplB-treated *Rag2^-/-^* mice were comparable to those observed in PBS-treated controls, indicating that cells of the adaptive immune system are the primary source of the type 2 cytokine response induced by SplB.

As protease activity is essential for many allergenic proteases to induce symptoms^30–32^, we next tested whether SplB’s catalytic activity was necessary to elicit AAI. We generated a catalytically inactive SplB mutant by introducing a point mutation (S193A) in its catalytic triad. The resulting SplB mut was completely devoid of protease activity in a specific cleavage assay (Fig. 1B), but retained a similar molecular structure compared to active SplB (Fig. E8 in the Online Repository). We then compared the effects of active SplB and SplB mut in our AAI model (Fig. 1A). Ovalbumin (OVA) i.t. was used as a specificity control. In this setting, we did not prime the animals with OVA i.p. and did not use any adjuvant. While the active SplB again induced a strong influx of eosinophils into the BALF and the lung tissue (Fig. 1C, D), OVA and the inactive SplB mutant failed to do so. Remarkably, there was only minimal recruitment of neutrophils to the BALF following sensitization and challenge with SplB (Fig. 1C). Eosinophil counts were almost three orders of magnitude higher than neutrophil counts. The number of neutrophils in the lungs did not change (Fig. 1E). Other lung cell populations are shown in figure E9 in the Online Repository. To investigate the systemic immune response, we analyzed the spleen and found a mild but significant eosinophilia in mice treated with the active SplB (Fig. 1F), suggesting that SplB-induced AAI causes systemic immune alterations. Next, we measured type 2 cytokines in the airways. Only active SplB elevated the IL-4-, IL-5-, and IL-13 concentrations in the BALF (Fig. 1G-I).

To evaluate the impact of SplB’s proteolytic activity on the humoral immune response, we measured total and antigen-specific IgE- and IgG levels in the serum (Fig. 2). Catalytic activity of SplB was necessary for production of SplB-specific IgE (Fig. 2A, B). However, SplB mut was able to provoke an antigen-specific IgG response, albeit less effectively than active SplB (Fig. 2E). These results demonstrate that the inactive SplB mutant is immunogenic but not allergenic. OVA-treated mice did not develop OVA-specific antibodies (Fig. 2C, F). Analysis of SplB-specific IgG subclasses showed an IgG1-dominated humoral response following exposure to SplB, whereas other IgG subclasses remained low (Fig. 2G-J). Histological examination of the lungs revealed that mice exposed to enzymatically active SplB had significantly thicker epithelium in their large airways than those treated with OVA or the inactive SplB mut (Fig. 3A). Mice treated with active SplB also showed increased mucus production (Fig. 3B), fibrosis (Fig. 3C) and eosinophil numbers in their lungs (Fig. 3D).

**Figure 2.**
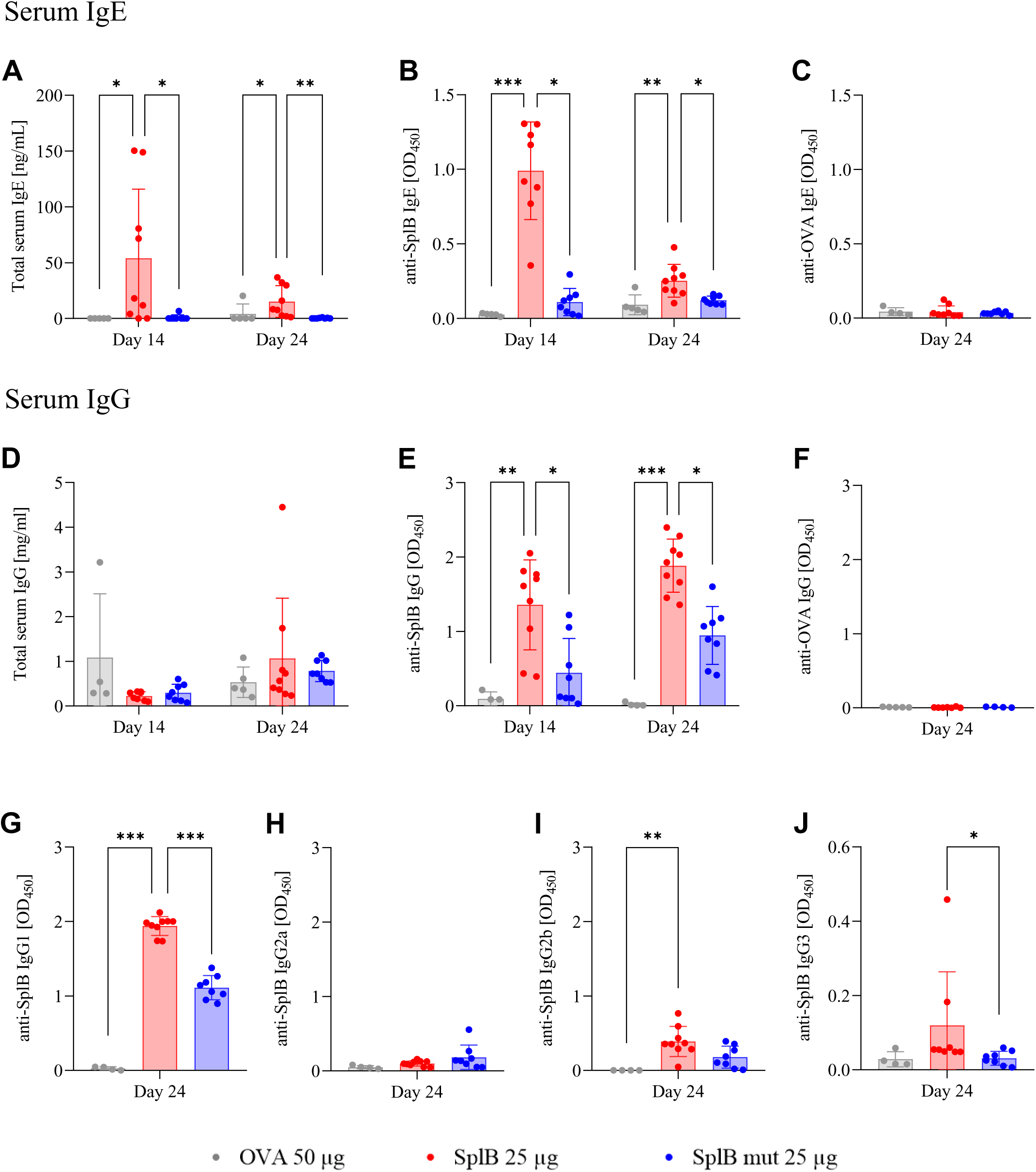
Catalytic activity of SplB is essential for IgE-, but not IgG secretion. Mice were treated with four doses of active SplB, inactive SplB mut, or OVA as control as shown in fig. 1A. Serum immunoglobulins were measured by ELISA on days 14 and 21 of the experiment. **A-C**, Serum IgE; total IgE (A), SplB-specific IgE (B) and OVA-specific IgE (C). **D-F**, Serum IgG. Total IgG (D), SplB-specific IgG (E) and OVA-specific IgG (F). **G**-**J**, Serum IgG subclasses. SplB-specific IgG1 (G), IgG2a (H), IgG2b (I), and IgG3 (J). n = 4-9. Results are presented as mean ±SD. *p < .05, **p < .01 and ***p < .005. Significance was determined using a Kruskal–Wallis test followed by Dunn’s multiple comparisons test.

**Figure 3.**
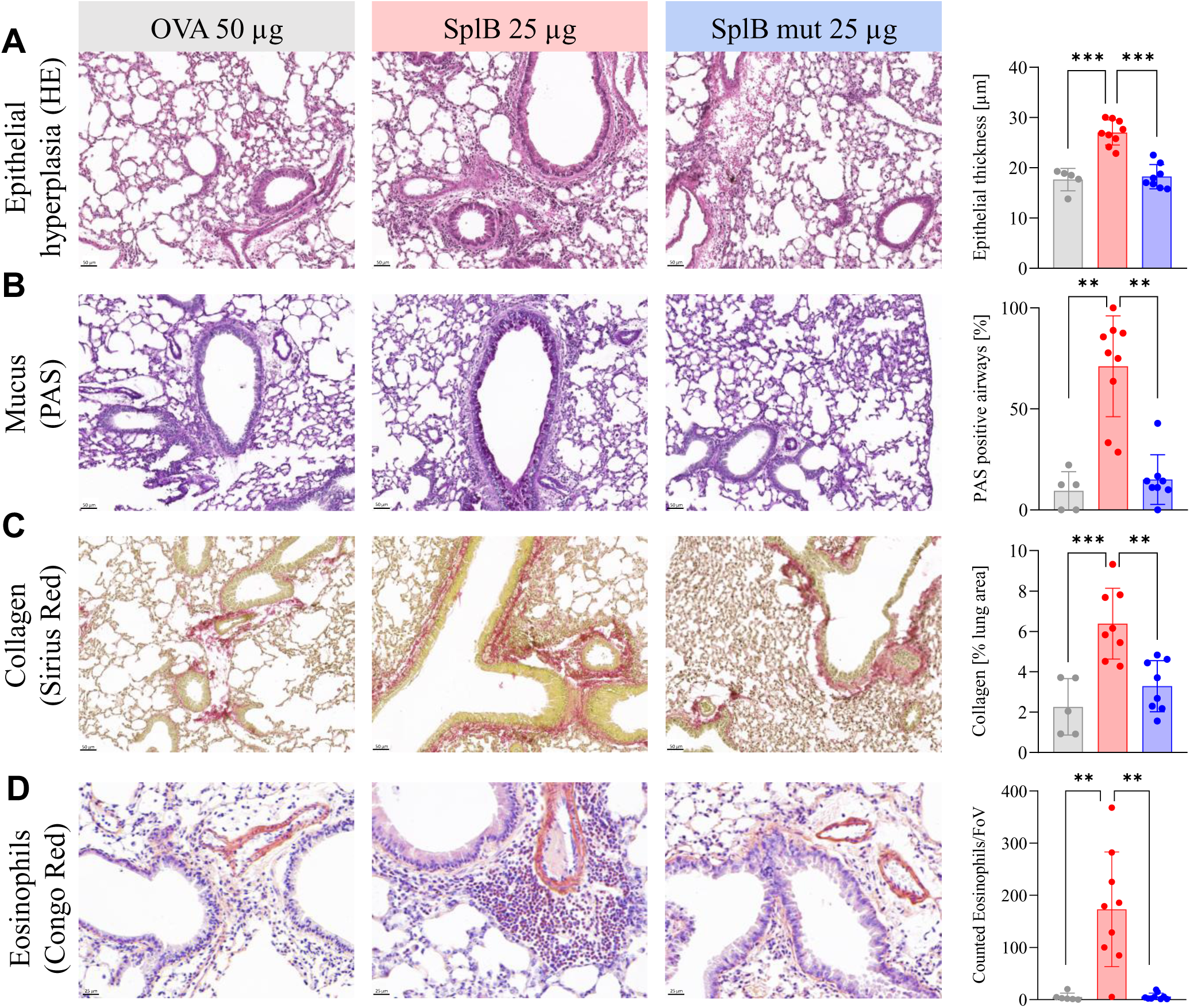
Only catalytically active SplB induces tissue remodeling and epithelial damage in the lungs. **A**, Hematoxylin and eosin (HE) staining of lung sections and thickness of airway epithelium. Epithelial thickness was determined by measuring the distance from the basement membrane to the apical epithelial surface on five points in three airways per section. **B**, Periodic acid-Schiff (PAS) staining for mucus in lung sections and percentage of PAS-positive airways. The percentage of PAS-positive airways was defined as the ratio of PAS-positive airways to the total number of airways per section. **C**, Sirius red staining for collagen in lung sections and percentage of the collagen-stained area in the total lung area. The percentage of collagen staining was calculated as the Sirius red-positive area divided by the total lung area. **D**, Congo red staining for eosinophils in lung sections and mean number of eosinophils per field of view. The mean number of eosinophils was calculated by counting the eosinophils in three different fields of view per lung section. n = 5-9. Results are presented as mean ±SD. *p < .05, **p < .01 and ***p < .005. Significance was determined using a one-way ANOVA followed by Tukey’s multiple comparisons test, except for B, where significance was determined using a Kruskal Wallis test followed by Dunn’s multiple comparisons test because the data were not normal distributed.

Treatment with active SplB significantly increased the albumin concentrations in the BALF, indicating impaired epithelial integrity (Fig. 4A). Additionally, IL-33 levels were increased in the lungs of mice exposed to active SplB (Fig 4B). Concentrations of thymic stromal lymphopoietin (TSLP), granulocyte-macrophage colony-stimulating factor (GM-CSF), IL-1a and IL-25 were below the detection limit.

**Figure 4.**
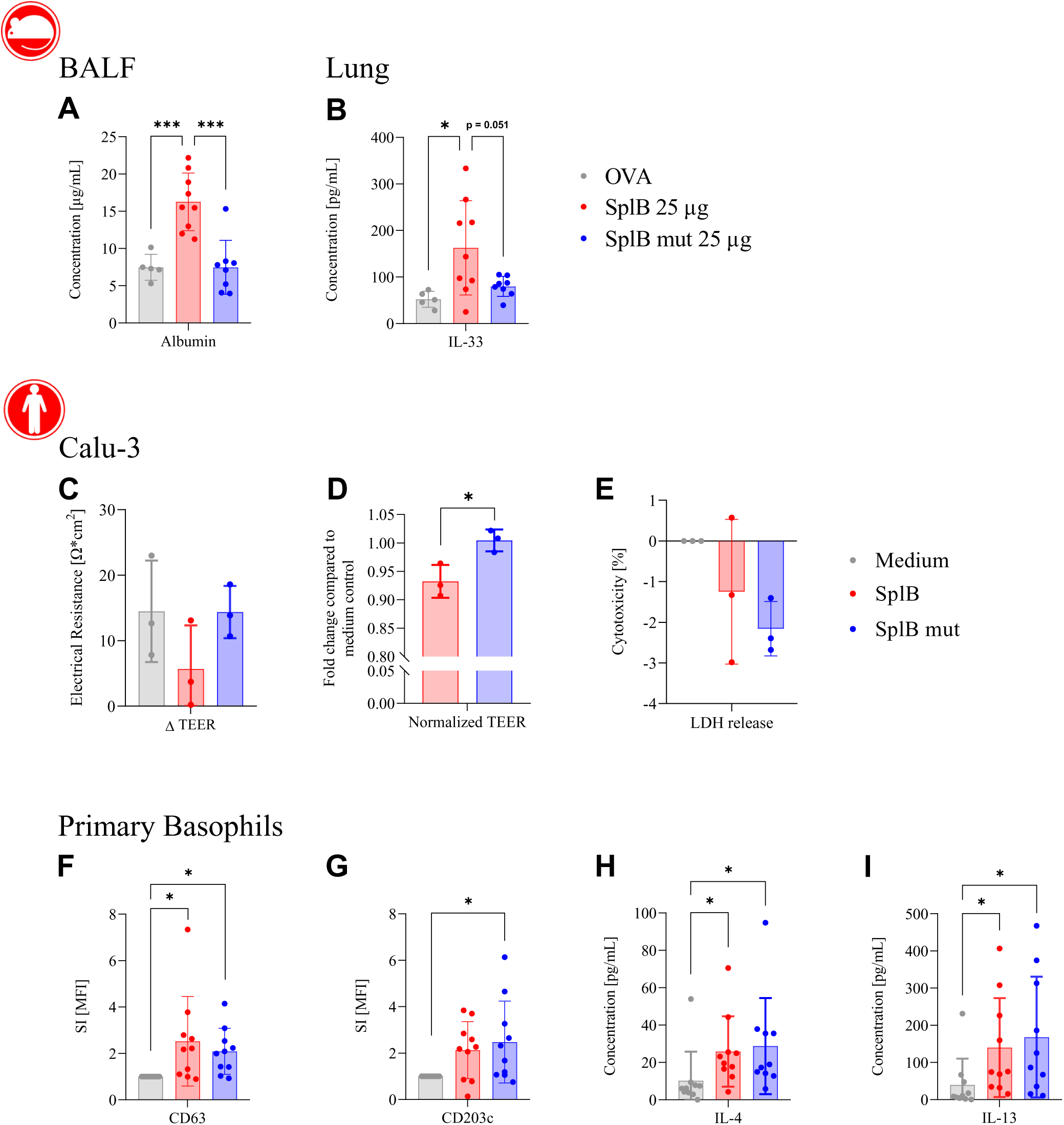
Active SplB damages the epithelial barrier and activates human basophils in vitro. **A** and **B,** mouse model, **C-I,** in vitro studies with human cells. **A**, Serum albumin concentration in the BALF measured by ELISA. **B**, IL-33 concentration was measured by ELISA in 50 µL buffer containing 50 µg of lung protein, n = 5-9, **C** and **D**, integrity of human Calu-3 airway epithelial cell layers exposed for 24 h to 20 µg/mL active SplB, inactive SplB mutant, or medium as a control. Barrier integrity was quantified by transepithelial electrical resistance (TEER). Changes in TEER were calculated as the difference between values after and before stimulation (C) and post-stimulation TEER values were normalized to medium-treated controls and expressed as fold change (D). **E,** cell death in Calu-3 cells exposed to 20 µg/mL of active SplB, SplB mutant, or medium quantified by LDH release. Data are shown as mean ± SD of three independent experiments, with each experiment representing the mean of technical triplicates. **F** and **G**, expression of the degranulation marker CD63 (F) and the activation marker CD203c (G) on primary basophils measured by flow cytometry. The stimulation index (SI) represents the fold change in marker expression relative to the unstimulated control. **H** and **I**, concentration of IL-4 (H) and IL-13 (I) in cell supernatants of primary basophils measured by CBA. n = 10. Results are presented as mean ± SD. *p < .05 and **p < .01. Significance was determined using a one-way ANOVA (A and B) followed by Tukey’s multiple comparisons test or Kruskal Wallis test followed by Dunn’s multiple comparisons test, depending on normal distribution of data.

Then, we turned to human airway epithelial cells to assess the relevance of our observations to the human situation. In order to test whether SplB can directly disturb the human epithelial barrier, we performed transwell experiments using Calu-3 epithelial monolayers. Treatment with active but not inactive SplB resulted in reduced transepithelial electrical resistance (TEER) demonstrating that the proteolytic activity of SplB compromises epithelial barrier integrity (Fig. 4C, D). To determine whether SplB-mediated barrier disruption was associated with epithelial cell death, LDH release was quantified after the exposure of the Calu-3 cells (Fig. 4E). SplB was not cytotoxic, showing that the increased permeability caused by enzymatically active SplB was not due to cell death. To further assess translational relevance of our findings, we studied the response of primary human basophils to SplB stimulation. The basophils were isolated from the blood of healthy volunteers with SplB-specific serum IgE. Exposure to SplB increased the expression of activation- and degranulation markers, and it significantly enhanced IL-4 and IL-13 secretion compared with medium-treated controls (Fig. 4F-I). Interestingly, the basophils responded to both the active SplB and the enzymatically inactive mutant, suggesting that while the protease activity of SplB is required for epithelial barrier disruption during the sensitization phase, structural recognition of the protein is sufficient to trigger effector responses.

Taken together, we found that only mice sensitized with enzymatically active SplB developed symptoms of allergic asthma and signs of a type 2 immune response, demonstrating that the enzymatic activity of SplB is necessary for the establishment of AAI.

### IL-33 is a crucial mediator in SplB-induced AAI

Asthma development often involves the alarmin IL-33^33^. To elucidate its role in SplB-induced allergic asthma, we co-administered the IL-33 inhibitor HpARI2 with SplB. In a parallel approach we tested the effect of SplB on *Il33^-/-^* mice. HpARI2 completely abolished the SplB-induced AHR following methacholine challenge (Fig. 5A) as well as the eosinophilic lung inflammation (Fig. 5B, C). In line with this, only WT mice but not *Il33^-/-^* mice showed a marked accumulation of eosinophils in the lung when challenged with SplB (Fig. 5B, C). In contrast, the anti-SplB IgE titers were increased in the sera of all SplB-treated mice, independent of the IL-33 activity (Fig. 5D). IL-33 was required for epithelial thickening and increased mucus production (Fig. 5E, F). Clearly, IL-33 is crucial for recruiting eosinophils to the site of inflammation, but it is not required for the immunoglobulin class switch in SplB-specific B cells.

**Figure 5.**
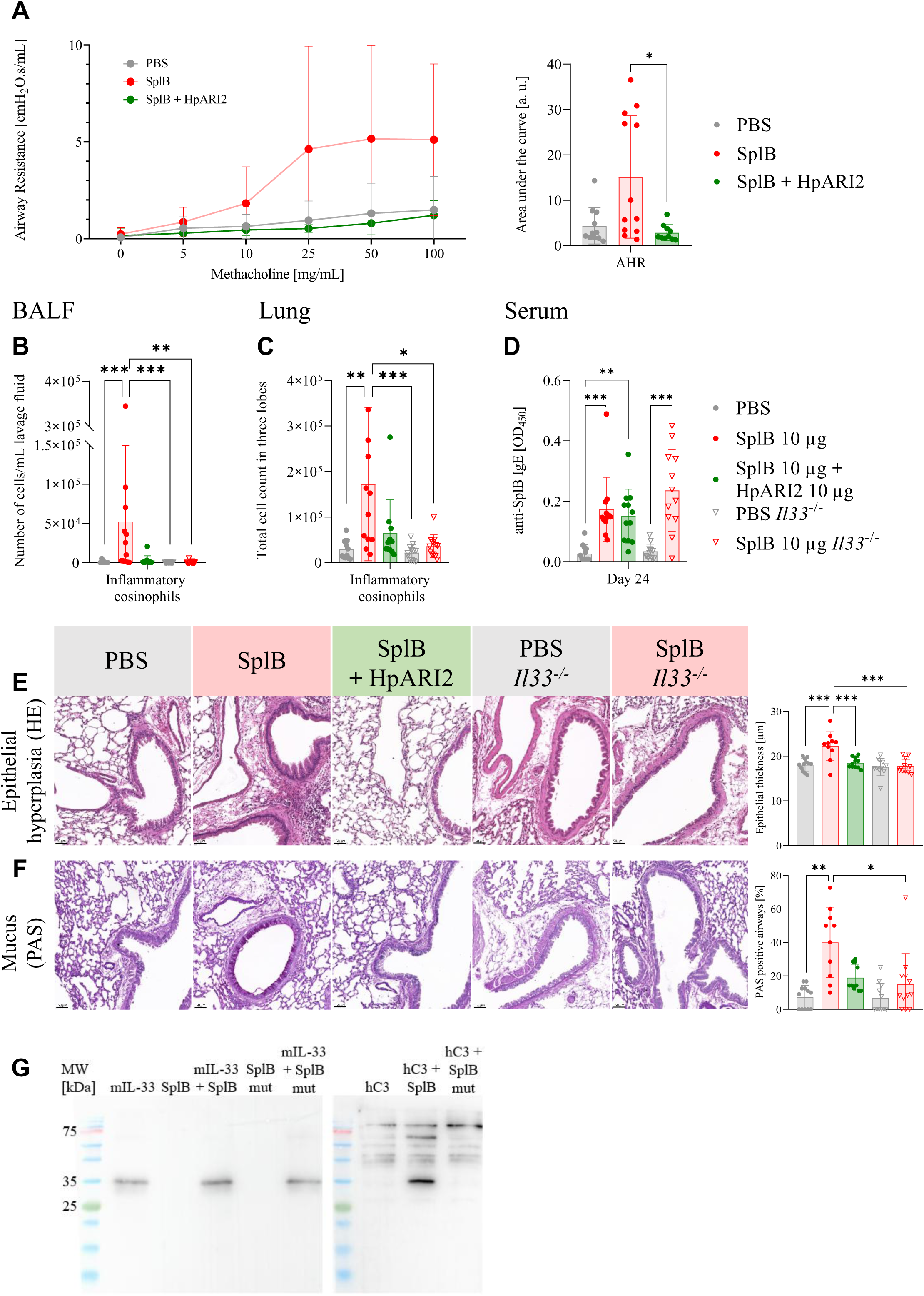
SplB-induced AAI is IL-33-dependent. WT C57BL/6J mice were treated with SplB or SplB + HpARI2, an IL-33 inhibitor. PBS-treated mice served as control. Additionally, C57BL/6J *Il33*^-/-^ mice were treated with either PBS or SplB. The treatment protocol is shown in fig. 1A. **A**, Airway resistance in response to increasing doses of methacholine was measured 72 hours after the last i.t. challenge using the forced oscillation technique. **B** and **C**, Numbers of inflammatory eosinophils in the BALF (B) and lung tissue (C) measured by flow cytometry. **D**, SplB-specific serum IgE on day 24 measured by ELISA. **E**, Epithelial thickness of airway epithelium and **F**, Mucus in lung sections (PAS staining) were determined as described in the legend of fig. 3. **G**, Enzymatic cleavage assays. Murine IL-33 or human complement factor C3 were incubated with active SplB or the inactive SplB mut for 24 h at 37 °C. Cleavage was visualized by Western blotting. n = 11-12. Results are presented as mean ± SD. *p < .05, **p < .01 and ***p < .005. Significance was determined using a Kruskal-Wallis test, except for E, where significance was determined using a one-way ANOVA followed by Tukey’s multiple comparisons test as data had normal distribution. AHR measurements were analyzed using a two-way ANOVA followed by Tukey’s multiple comparisons test.

Numerous allergens cleave the alarmin IL-33, which increases its biological activity and promotes the activation of innate immune cells and Th2 cells^18^. We therefore asked whether SplB targets IL-33 and incubated full-length murine IL-33 (mIL-33) with active SplB. Immunoblotting revealed that SplB did not cleave mIL-33, although it cleaved the human complement factor C3 under the same conditions (Fig. 5G and published data^34^). Therefore, mIL-33 is not a substrate of SplB.

### PAR2 is dispensable for the development of SplB-induced AAI

PAR2 activation by protease allergens promotes type 2 airway inflammation and asthma severity is correlated with increased PAR2 expression^35^. Therefore, we examined the role of PAR2 in SplB-induced murine AAI. To this end, we treated *F2rl1^-/-^* mice with SplB or applied a PAR2-blocking monoclonal antibody (SAM-11) to WT mice. *F2rl1^-/-^* mice exposed to SplB displayed the same number of inflammatory cells in their BALF and lungs as their WT counterparts (Fig. 6A, B). Likewise, WT and *F2rl1^-/-^* mice had similar titers of anti-SplB IgE in their sera (Fig. 6C). In line with this, PAR2 deficiency had no impact on epithelial thickness or mucus production (Fig. 6D, E). However, remarkably, it significantly reduced lung fibrosis (Fig. 6F). Similarly, blocking of PAR2 with the monoclonal antibody SAM-11 failed to protect mice from AAI, but reduced lung fibrosis (Fig. E10 in the Online Repository).

**Figure 6.**
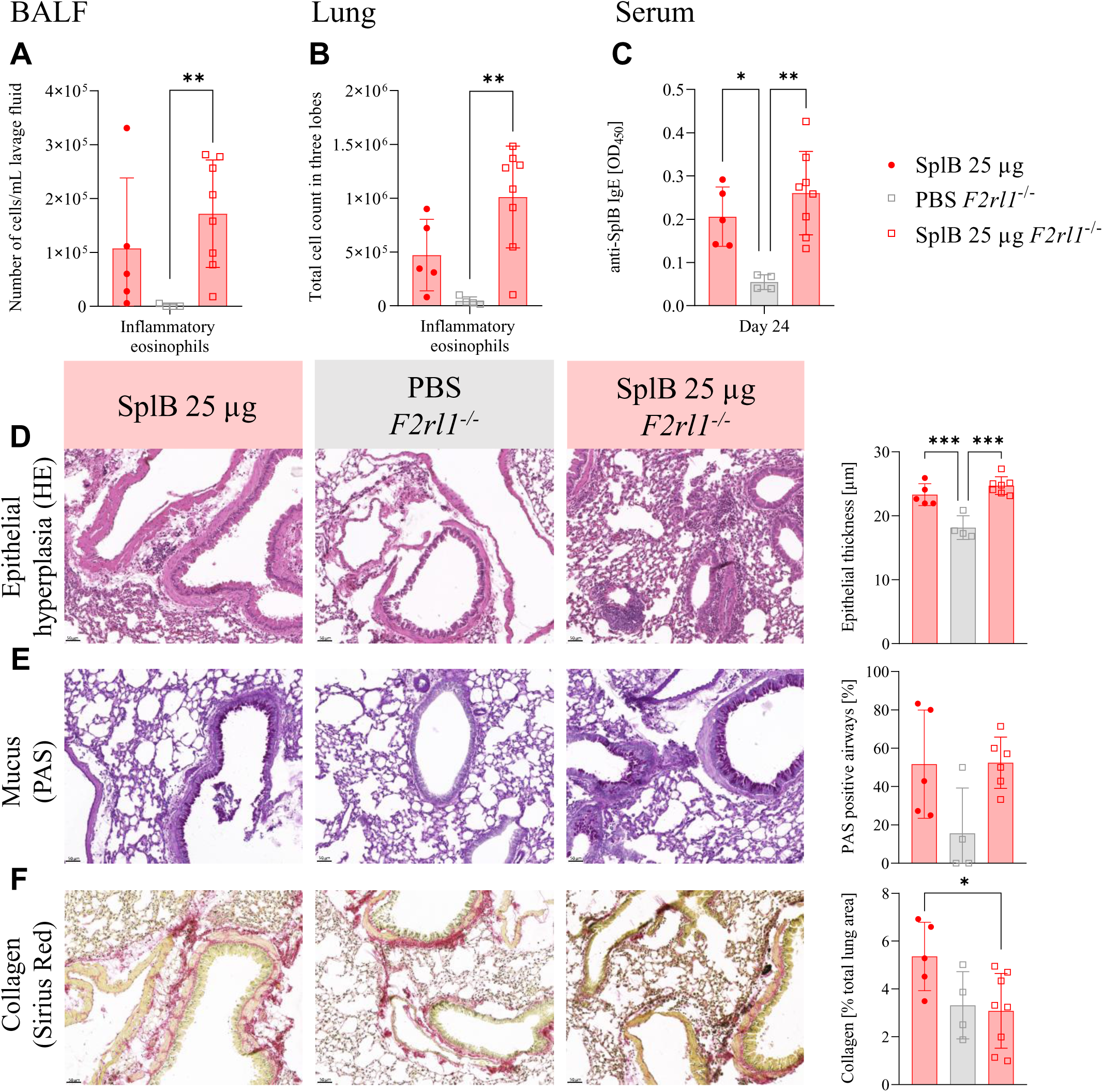
SplB-induced AAI is independent of PAR2 activation. WT or *F2rl1^-/-^* (PAR2-deficient) C57BL/6J mice were treated with SplB; PBS-treated *F2rl1^-/-^* mice served as controls. **A** and **B**, Numbers of inflammatory eosinophils in the BALF (A) and lung tissue (B) measured by flow cytometry. **C**, SplB-specific serum IgE measured by ELISA. **D**, Epithelial thickness of airway epithelium, **E**, Mucus in lung sections (PAS staining) and **F**, Collagen (Sirius Red) were determined as described in the legend of fig. 3. n = 4-8. Results are presented as mean ±SD. **p < .01. Significance was determined using a Kruskal-Wallis test followed by Dunn’s multiple comparisons test (A-C) or one-way ANOVA followed by Tukey’s multiple comparisons test (D-F), depending on normal distribution of data.

## Discussion

In this study, we show that the protease SplB, which is secreted by *S. aureus*, is a potent allergen that induces type 2 airway inflammation in mice. Four intratracheal applications were sufficient to induce AAI without the need for adjuvants or prior intraperitoneal priming. The AAI phenotype induced by SplB was characterized by increased AHR, a massive influx of eosinophils into the airways and the lung, lung fibrosis, and the production of type 2 cytokines as well as of SplB-specific IgE. Various factors contributed to murine AAI development in distinctive ways: SplB’s enzymatic activity was essential for eosinophilic airway inflammation, fibrosis and IgE induction, but not for the production of specific IgG. IL-33 was necessary for eosinophil infiltration and fibrosis, but not for the production of specific IgE or IgG. PAR2 was involved in lung fibrosis but not the other symptoms of AAI.

In our AAI model, the allergenic potential of SplB was strictly dependent on its proteolytic activity, as a catalytically inactive SplB mutant did not elicit AAI nor SplB-specific IgE. Nevertheless, the inactive SplB mutant, but not OVA, induced a specific IgG response. Clearly, enzymatically inactive SplB is immunogenic but not allergenic. The absence of an antibody response to OVA might be attributed to immune tolerance due to OVA’s eukaryotic origin and structural similarities with host proteins^36,37^. In contrast, the SplB mutant, which was genetically inactivated by a single point mutation, appears to be structurally very similar to the active bacterial enzyme. Thus, the distinctive immune response to the active and inactive bacterial protease suggests that, in its natural form, SplB has a dual effect on the immune system: (i) allergenicity, i.e., type 2 polarization requiring protease activity and (ii) immunogenicity facilitated by structural motifs, possibly conserved in bacteria, that are recognized by innate immune cell receptors^38^.

Since catalytically active SplB increased IL-33 concentrations in the lung, we hypothesized that the protease induces AAI via the IL-33 signaling pathway. This was supported by the results of two experimental strategies, IL-33 blockade by HpARI2, a potent anti-IL-33 agent originally derived from helminths^39^, and the use of IL-33-deficient animals. The IL-33 function was necessary for the eosinophilic lung inflammation but not for the generation of SplB-specific IgE. We conclude that mediators other than IL-33 must be responsible for the sensitization to SplB, which involves the maturation of immature DCs into cDC2s that drive the polarization of Th2 cells^40,41^. A candidate for this is the alarmin TSLP known to be essential for cDC2 maturation^42^. The concentrations of TSLP, however, remained below the limit of detection in the airways and lungs of the SplB-treated mice. Later during sensitization, T follicular helper (Tfh) cells are critical for B cell activation and the immunoglobulin class switch to IgE in the germinal centers of the secondary lymphoid organs^43^. Tfh cells, however, do not express the IL-33 receptor ST2 and are thus independent of IL-33 signaling^44^. The absence of eosinophilic lung inflammation in mice without active IL-33 could be explained by the role of mast cells. Mast cells express ST2 at high density, and IL-33 lowers their degranulation threshold^45,46^. Degranulating mast cells release large amounts of IL-5, which is critical for eosinophil recruitment^47^. In addition, group 2 innate lymphoid cells (ILC2s) represent a major source of IL-5, and their cytokine production is strongly dependent on IL-33^48^. However, ILC2 activity alone cannot account for the observed phenotype, as IL-33–competent *Rag2*-deficient mice failed to develop asthmatic features. Thus, the adaptive immune system as well as catalytically active SplB and IL-33 were necessary for AAI development. However, there is no direct link between the latter as the protease SplB does not target murine IL-33.

The translational relevance of our findings should be considered in light of known species-specific properties of the IL-33/ST2 axis. While murine and human IL-33 only share around 55% sequence homology, the overall architecture and function of the IL-33/ST2 pathway are highly conserved^33,49^. Notably, the main cells that produce IL-33 in the lungs differ between species: alveolar type II epithelial cells are the main source of the alarmin in mice, whereas bronchial epithelial cells are the main producers in humans^50,51^. Additionally, variations in IL-33 expression patterns and proteolytic processing have been documented^52^. While these factors may influence the magnitude or localization of IL-33-driven responses, there is extensive experimental and clinical evidence supporting a central role for the IL-33/ST2 pathway in human asthma^33^. Therefore, despite these differences, the IL-33 dependence observed in our model probably reflects a biologically relevant mechanism in the human disease.

We reasoned that SplB may cause allergic asthma symptoms in mice by cleaving a substrate other than IL-33, possibly the membrane-bound protease sensor PAR2, which is targeted by several known protease allergens^16,53^. However, most symptoms in SplB-mediated AAI were independent of PAR2. This is not unprecedented, as other studies have shown that, for example, HDM can also induce AAI independent of PAR2^54,55^. Notably, the absence of PAR2 significantly reduced the extent of lung fibrosis, corroborating previous observations in a murine model of HDM-induced asthma^54,56^. While PAR2 is the best-characterized protease-activated receptor in allergic airway inflammation, additional mechanisms of protease sensing may be involved. The widely expressed protease sensor PAR4 can also be activated by allergen-derived proteases, resulting in intracellular calcium signaling that can promote epithelial activation and the release of alarmins such as IL-33 and TSLP^57,58^. We cannot exclude that PAR4 partially compensates for the absence of PAR2 in our model and contributes to the observed inflammatory response. Our transwell experiments with human airway epithelial cells show that enzymatically active SplB directly disturbed the epithelial barrier, indicating that protease-mediated epithelial damage itself is a major driver of the observed phenotype. We therefore favor a scenario in which direct epithelial injury constitutes the primary mechanism underlying SplB-induced airway inflammation, whereas activation of PAR family members, including PAR4, may represent an additional contributory pathway.

Epithelial barriers are the first line of defense against environmental threats like pathogens and allergens. *S. aureus* needs to overcome host barriers to persist and cause infections. A previous study has shown that Spls play a role in bacterial spread in a rabbit pneumonia model, suggesting that the bacterial proteases damage barriers^59^. Indeed, SplB can disrupt endothelial barriers and cause vascular leakage^60^. In our model of AAI, i.t. application of SplB impaired the integrity of the airway epithelial barrier, as demonstrated by increased leakage of serum albumin into the airways. We also observed disruption of barrier function by enzymatically active SplB in a transwell assay with human epithelial cells, suggesting that our finding is translatable to human airway epithelium. This functional impairment is consistent with a direct SplB attack on host barrier cells or junctional proteins. Previous studies have characterized the substrate specificity of SplB on human proteins, identifying complement factors and ubiquitin-like proteases as targets^34,61^. More recently, our group has identified further SplB substrates, namely intermediate filament proteins (desmin, vimentin, nestin), heat shock protein pi, a-enolase, and IgG^62^. This suggests that SplB targets multiple host proteins involved in immune regulation and cellular structural integrity, which may collectively contribute to epithelial barrier dysfunction. However, the initial molecular target(s) of SplB in the pathogenesis of murine AAI remain to be elucidated in future studies. This is the main limitation of our study.

The clinical relevance of SplB should be considered in the broader context of *S. aureus* airway colonization. In colonized individuals, SplB is encountered by the immune system together with numerous other staphylococcal virulence factors. Many of these, notably Hla, Plc, GlpQ and Lip, induce a type 1/ type 3 responses^26,63^ as they are required for immune protection against *S. aureus* infection^64–66^. The *S. aureus* superantigens SEA-SEE are exceptional as they can promote type 2 responses, including specific as well as polyclonal IgE production^67^. Thus, by disrupting the airway barrier, SplB could act in synergy with superantigens, amplifying a local type 2 immune response.

We have identified SplB as a potent bacterial allergen that causes eosinophilic lung inflammation on its own without exogenous adjuvants or systemic priming. SplB elicits the production of allergen-specific IgE in both mice and humans^25,26^, and promoted airway hyperresponsiveness. Other members of the Spl family, namely SplD and SplF, also induce AAI in mice^24^. These findings suggest that *S. aureus* may play a more active role in allergic asthma than previously thought and they advance our understanding of the role of *S. aureus* airway colonization in the pathogenesis of asthma. Two mutually non-exclusive hypotheses have been proposed: (1) Asthma and the associated type 2 immune response facilitate *S. aureus* colonization and (2) *S. aureus* colonization creates a type 2 microenvironment promoting allergic sensitization towards harmless antigens^9^. Based on our results we propose a third hypothesis to complement these two scenarios: (3) *S. aureus* secretes allergens that can induce allergic asthma development and trigger asthma attacks.

To date, only one bacterial allergen is listed in the WHO/IUIS Allergen Nomenclature Sub-Committee’s database of allergenic molecules, the serine protease nattokinase from *Bacillus subtilis var. natto* [www.allergen.org]^68,69^. Our findings suggest that we may have underestimated the role that bacterial allergens play in disease development, and they encourage the scientific community to increase its efforts to study and clarify this issue.

## Acknowledgements

We thank Dr. Sabine Berg, Sabine Prettin and Shruthi Peringathara for their support with the animal experiments and Katrin Schoknecht and Fawaz Al-Sholui for technical assistance. We are grateful that Deutsche Forschungsgemeinschaft (DFG, German Research Foundation) is funding our research (project number 443535983).

## Declaration of generative AI and AI-assisted technologies in the manuscript preparation process

During the preparation of this work the authors used the platform DeepL write for text revision. After using this tool, the authors reviewed and edited the content as needed and they take full responsibility for the content of the published article.

## Abbreviations

AAI: Allergic airway inflammation
AHR: Airway hyperreactivity
AMC: Amino-methyl-cumarin
APC: Allophycocyanin
BALF: Bronchoalveolar lavage fluid
BCA: Bicinchoninic acid
CBA: Cytometric bead array
CD: Cluster of differentiation
DC: Dendritic cells
ELISA: Enzyme-linked Immunosorbent Assay
FCS: Fetal calf serum
FITC: Fluorescein isothiocyanate
GlpQ: Glycerophosphoryl diester phosphodiesterase
GM-CSF: Granulocyte-macrophage colony-stimulating factor
HDM: House dust mite
HE: Hematoxylin Eosin
Hla: a-Hemolysin
HpARI2: Heligmosomoides polygyrus Alarmin Release Inhibitor 2
HRP: Horseradish peroxidase
Ig: Immunoglobulin
IL: Interleukin
ILC2: Type 2 innate lymphoid cell
i.t.: intratracheally
Lip: Lipase
MHC: Major histocompatibility complex
OD: Optical density
OVA: Ovalbumin
PAR: Protease-activated receptor
PAS: Periodic acid–Schiff
PBS: Phosphate-buffered saline
PE: Phycoerythrin
PerCP: Peridinin-chlorophyll-protein complex
Plc: Phospholipase C
Rag2: Recombination-activating gene
RT: Room temperature
SDS-PAGE: Sodium dodecyl sulfate polyacrylamide gel electrophoresis
Spl: Serine protease–like protein
RIPA: Radioimmunoprecipitation Assay
Tfh: T follicular helper
Th2: T helper cell type 2
TSLP: Thymic stromal lymphopoietin
WT: Wild type

---

**Figure E1.** Protein sequences of the recombinant preparations of active SplB and catalytically inactive SplB mut. The SplB sequence is derived from the *S. aureus* strain USA300_FPR3757. The expressed SplB and SplB mut proteins contain a signal peptide (grey), which is cleaved off upon secretion by *S. aureus* RN4220, which served as expression system. In the SplB mut protein the catalytic serine was switched to an alanine ((S193A, yellow). A C-terminal Twin-Strep-tag® used for purification is highlighted in blue. The amino acid linker between the SplB protein and the Twin-Strep-tag®is marked in green. Tag and linker were not removed from the mature SplB resp. SplB mut proteins.

**Figure E2.** Gating strategy for immune cell phenotyping in the BALF. BALF was collected and centrifuged, the supernatant was removed and the cells were stained for flow cytometry analysis. **A-C** Gating of single cells, exclusion of debris and duplets. **D**, Gating of living cells. **E**, Distinguishing inflammatory eosinophils (SiglecF^+^; CD11c^-^) and alveolar macrophages (SiglecF^+^; CD11^+^) from SiglecF^-^ cells. **F**, The SiglecF^-^ cells are further distinguished in neutrophils (SiglecF^-^; Ly6G^+^) and CD4^+^ T cells (SiglecF^-^, CD4^+^). The following antibodies were used: anti-mouse CD4-VioBlue (130-118-696), CD11c-allophycocyanin (APC) (130-110-839), Ly6-G-fluorescein isothiocyanate (FITC) (130-120-820) and Siglec-F-phycoerythrin (PE)-Vio770 (130-112-334).

**Figure E3.** Gating strategy for DC subpopulations and eosinophils from lung tissue. Three lobes of the right lung (superior, middle and inferior lobe) were digested and stained for flow cytometry analysis. **A-C**, Gating of single cells, exclusion of debris and duplets. **D**, Gating of living cells. **E**, Distinguishing inflammatory eosinophils (SiglecF^+^; CD11c^-^), resident eosinophils (SiglecF^+^ low; CD11c^+^), alveolar macrophages (SiglecF^+^; CD11^+^) from SiglecF^-^ cells. **F**, The SiglecF^-^ cells are further distinguished in dendritic cells (SiglecF^-^; MHCII^+^; CD11c^+^) and MHCII^-^ cells. **G**, MHCII^-^ cells were separated into Ly6G^-^ cells and neutrophils (Ly6G^+^). **H-I**, DCs subpopulations were further distinguished into type 1 cDCs (SiglecF^-^; MHCII^+^; CD11c^+^, CD11b^-^, CD103^+^), type 2 cDCs (SiglecF^-^; MHCII^+^; CD11c^+^, CD11b^+^, CD103^-^, CD64^-^) and moDCs (SiglecF^-^; MHCII^+^; CD11c^+^, CD11b^+^, CD103^-^, CD64^+^). The following antibodies were used: anti-mouse Ly6-G-FITC (130-120-820), MHC Class II-VioGreen (130-112-395), CD103-APC (130-111-686), CD11b-PE-Vio770 (130-113-808), CD64-PE (130-118-684), Siglec-F-PE-Vio615 (130-112-330) and CD11c-VioBlue (130-110-843). The following antibodies were used for dump channel: anti-mouse CD19-APC-Vio770 (130-112-038), CD3-APC-Vio770 (130-119-793) and CD49b-APC-Vio770 (130-116-435).

**Figure E4.** Gating strategy for T cell subpopulations from lung tissue. Three lobes of the right lung (superior, middle and inferior lobe) were digested and stained for flow cytometry analysis. **A-C**, Gating of single cells, exclusion of debris and duplets. **D**, Gating of living cells. **E**, CD3^-^ cells were separated into NK1.1^-^ cells and NK cells (CD3^-^, NK1.1^+^). **F**, CD4^-^ T cells (CD3^+^, CD4^-^), CD4^+^ T cells (CD3^+^, CD4^+^) were distinguished from CD3^-^ cells. **G**, CD4^-^ T cell subpopulations were distinguished into naïve T cells (CD3^+^, CD4^-^, CD62L^+^, CD44^-^), effector T cells (CD3^+^, CD4^-^, CD62L^-^, CD44^+^) and memory T cells (CD3^+^, CD4^-^, CD62L^+^, CD44^+^). **H**, CD4^+^ T cell subpopulations were distinguished into naïve T cells (CD3^+^, CD4^+^, CD62L^+^, CD44^-^) and effector T cells (CD3^+^, CD4^+^, CD62L^-^, CD44^+^). The following antibodies were used: anti-mouse CD3-peridinin-chlorophyll-protein complex (PerCP)-Vio700 (130-120-826), MHC Class II-VioGreen (130-112-395), CD4-Brilliant Violet 605 (BioLegend^®^, San Diego, USA; 100548), CD11b-PE-Vio770 (130-113-808), CD44-VioBright FITC (130-120287), NK1.1-PE (130-120-506), CD11c-VioBlue (130-110-843) and CD62L-APC (130-112-837).

**Figure E5.** Gating strategy for immune cells from the spleen. A single cell suspension was prepared rom a part of the spleen and stained for flow cytometry. **A** and **B**, Gating of single cells, exclusion of debris and duplets. **C**, Distinguishing inflammatory eosinophils (SiglecF^+^, CD11c^-^) from SiglecF^-^ cells. **D**, CD3^+^ cells were distinguished from CD3^-^ cells. **E**, B-cells (CD3^-^, CD19^+^, B220^+^) were distinguished. **F**, Distinguishing Th cells (CD3^-^, CD4^+^, CD8^-^) from cytotoxic T cells (CD3^-^, CD4^-^, CD8^+^). The following antibodies were used: anti-mouse Siglec-F-PE-Vio770 (130-112-334), CD11c-APC (130-110-839), CD3-PerCP-Vio700 (130-120-826), CD4-VioBlue (130-118-696), CD8-PE (BioLegend^®^, San Diego, USA; 100708), CD45R-Brilliant Violet 510 (BioLegend^®^, San Diego, USA; 103248) and CD19-Brilliant Violet 650 (BioLegend^®^, San Diego, USA; 115541).

**Figure E6.** SplB induces allergic airway inflammation in mice at different concentrations. **A**, Airway resistance in response to increasing doses of methacholine was measured 72 hours after the last i.t. challenge using a forced oscillation technique. **B** and **C**, Numbers of inflammatory eosinophils in the BALF (B) and lung tissue (C) measured by flow cytometry. **D**, SplB-specific serum IgE measured by ELISA. **E**, Epithelial thickness of airway epithelium and **F**, Mucus in lung sections (PAS staining) were measured as described in the legend of fig. 3. n = 1-7. Results are presented as median ± interquartile range. *p < .05, **p < .01 and ***p < .005. Significance was determined using a Kruskal–Wallis test followed by Dunn’s multiple comparisons test., except for E, where significance was determined using a one-way ANOVA followed by Tukey’s multiple comparisons test because the data showed normal distribution. AHR measurements were analyzed using a two-way ANOVA followed by Tukey’s multiple comparisons test.

**Figure E7.** AAI induction by SplB requires an intact adaptive immune system. WT or *Rag2*^-/-^ C57BL/6J mice were treated with SplB as shown in fig. 1 in the main paper, PBS-treated *Rag2^-/-^* mice served as control. **A** and **B**, Numbers of inflammatory eosinophils (A) and neutrophils (B) in the BALF measured by flow cytometry. **C**, Number of inflammatory eosinophils in the lung tissue measured by flow cytometry. **D** and **E**, Concentrations of IL-5 (D) and IL-13 (E) in the BALF measured by CBA. **F**, Epithelial thickness of airway epithelium and **G**, Mucus in lung sections (PAS staining) were measured as described in the legend of fig. 3. n = 3-6. *p < .05, **p < .01 and ***p < .005. Results are presented as mean ±SD. Significance was determined using a Kruskal–Wallis test followed by Dunn’s multiple comparisons test (A-C) or a one-way ANOVA followed by Tukey’s multiple comparisons test (F-G), depending on normal distribution of data.

**Figure E8.** Analytical size exclusion chromatography (SEC) of active SplB (named SplB wt in the figure) and the inactive SplB mutant. The data show that the monomeric forms (peak 3) of both SplB wt and SplB mutant run with an almost identical apparent molecular mass (Mr), indicating conservation of their structure. In the size exclusion chromatography, the native proteins form multimers (peaks 1 and 2) in different proportions (A, C), while the denaturing SDS-PAGE shows only the bands of the monomers (B). **A**, Overlay of size exclusion chromatography runs of active SplB (blue) and SplB mutant (pink). The analytical SEC runs were performed using a Superdex 75 GL 10/300 column. Protein was detected using the absorption at 280 nm. P1 = peak 1, P2 = peak 2, P3 = peak 3. Peak 3 corresponds to the monomeric form of SplB. **B**, SDS-PAGE analysis of the protein-containing SEC elution fractions of active SplB (left) and SplB mutant (right). M: protein standard PegGold. Left panel, active SplB: the SEC elution fraction 20 corresponds to P1 of the analytical SEC shown in (A), elution fractions 23 – 26 correspond to P2 of the SEC, elution fractions 27 – 30 correspond to P3. Right panel, SplB mutant: the SEC elution fraction 20 corresponds to P1 of the analytical SEC shown in (A); elution fractions 25 – 32 correspond to P3 of SEC. **C**, Calibration curve and corresponding fitting equation of the Superdex 75 GL 10/300 column calculated with the known molecular weights and the experimentally determined elution volumes of Aprotinin, Ribonuclease A, GFP, Ovalbumin and Conalbumin (right). Calculation of apparent molecular weights of different isoforms of SplB, which eluted at different volumes in the native analytical SEC. The ratio of apparent molecular weight and theoretical molecular weight corresponds to the mono- or oligomeric states of the protein.

**Figure E9.** Immune cell populations in the lungs of mice exposed to catalytically active or inactive SplB. Female WT C57BL/6J mice were treated with active SplB or the catalytically inactive SplB mut; OVA-treated mice served as control. Numbers of **A**, resident eosinophils; **B**, alveolar macrophages; **C**, natural killer (NK) cells; **D**, dendritic cells (DCs), **E**; DC subpopulations, and **F**, CD4^+^ T-cells. **G**, ratios of CD4^+^ T-cell subpopulations. **H**, numbers of CD4^-^ T-cells; and **I**, ratios of CD4^-^ T-cell subpopulations in the lung measured by flow cytometry. n = 5-9. Results are presented as mean ±SD. *p < .05 and **p < .01. Significance was determined using a Kruskal–Wallis test followed by Dunn’s multiple comparisons test.

**Figure E10.** SplB-induced AAI is independent of PAR2 function. WT C57BL/6J mice were treated with SplB or SplB + SAM-11, a PAR2-inhibiting monoclonal antibody. Mice treated with SplB + IgG2a served as isotype control. **A** and **B**, Numbers of inflammatory eosinophils in the BALF (A) and lung tissue (B) measured by flow cytometry. **C**, SplB-specific IgE concentrations measured by ELISA. **D**, Epithelial thickness of airway epithelium, **E**, Mucus in lung sections (PAS staining) and **F**, Collagen quantification (Sirius Red) were determined as described in the legend of fig. 3. n = 5. **p < .01. Results are presented as mean ±SD. Significance was determined using a Kruskal–Wallis test followed by Dunn’s multiple comparisons test.

