## Supplement Figure E1 for "The *Staphylococcus aureus* serine protease-like protein B is a potent allergen in a murine asthma model"

Protein sequence of SplB WT as expressed in *S. aureus* RN4220 pTripleTREP_splB.wt:

MNKNVVIKSLAALTILTSVTGIGTTLVEEVQQTAKAENNVTKVKDTNIFPYTGVVAF
KSATGFVVGKNTILTNKHVSKNYKVGDRITAHPNSDKGNGGIYSIKKIINYPGKEDVS
VIQVEERAIERGPKGFNFNDNVTPFKYAAGAKAGERIKVIGYPHPYKNKYVLYESTG
PVMSVEGSSIVYSAHTESGNSGSPVLNSNNELVGIHFASDVKNDDNRNAYGVYFTPE
IKKFIAENIDKGSWSHPQFEKGGGSGGGSGGSAWSHPQFEK

Protein sequence of SplB mutant as expressed in *S. aureus* RN4220 pTripleTREP_splB.mut:

MNKNVVIKSLAALTILTSVTGIGTTLVEEVQQTAKAENNVTKVKDTNIFPYTGVVAF
KSATGFVVGKNTILTNKHVSKNYKVGDRITAHPNSDKGNGGIYSIKKIINYPGKEDVS
VIQVEERAIERGPKGFNFNDNVTPFKYAAGAKAGERIKVIGYPHPYKNKYVLYESTG
PVMSVEGSSIVYSAHTESGNAGSPVLNSNNELVGIHFASDVKNDDNRNAYGVYFTPE
IKKFIAENIDKGSWSHPQFEKGGGSGGGSGGSAWSHPQFEK
