## Supplementary figures and images for "The *Staphylococcus aureus* serine protease-like protein B is a potent allergen in a murine asthma model"

### Supplement Figure E2

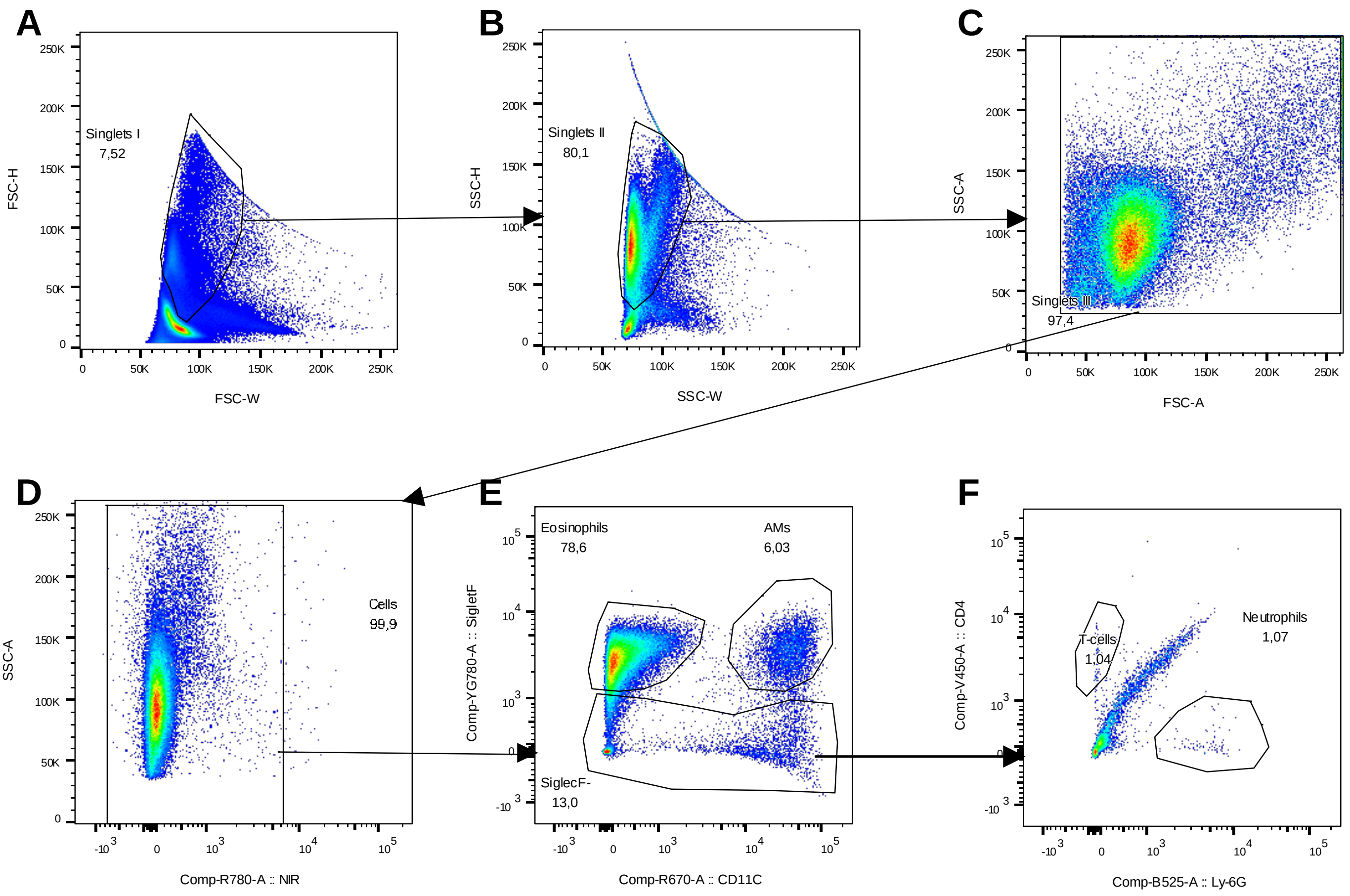

### Supplement Figure E3

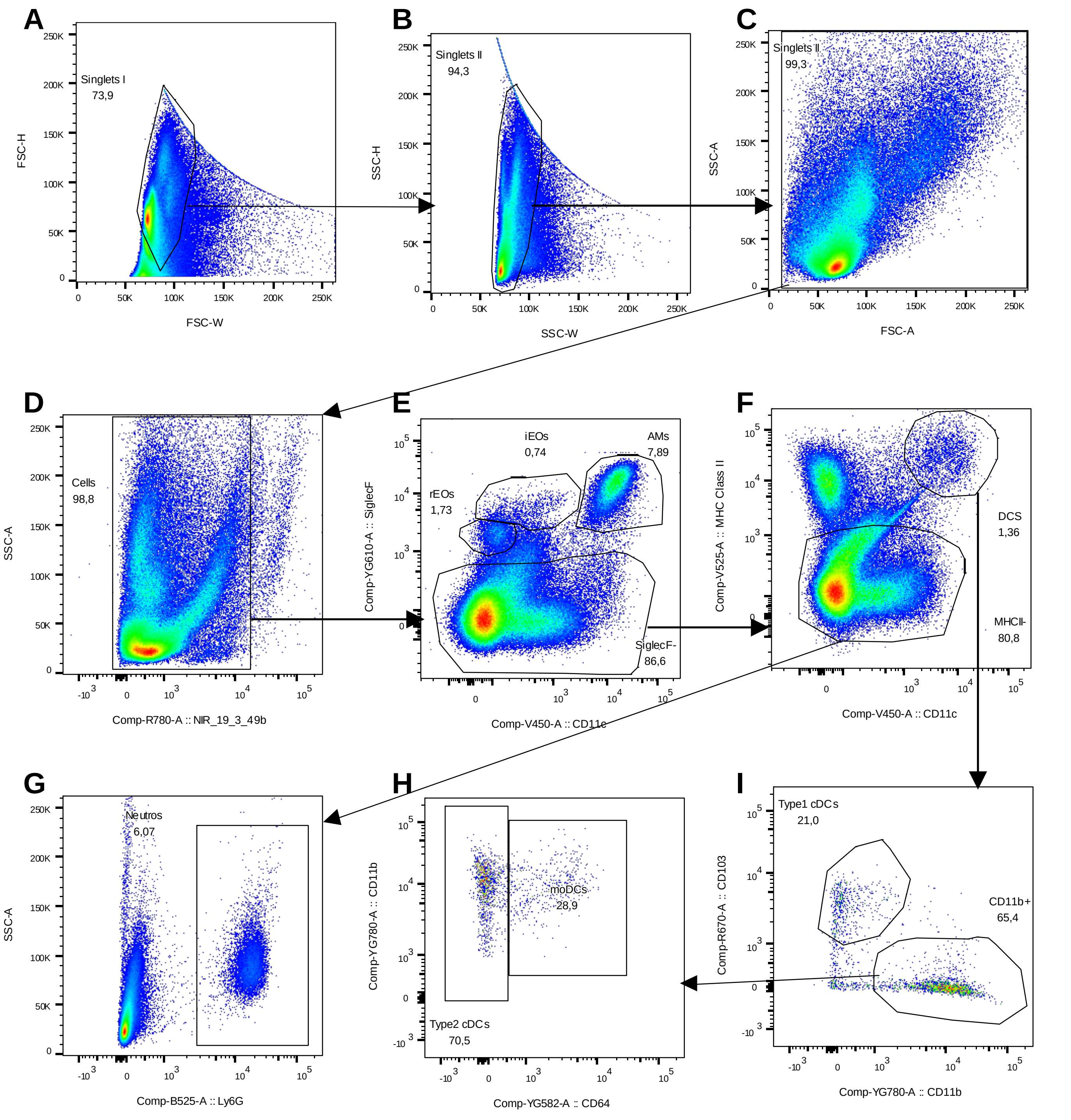

### Supplement Figure E4

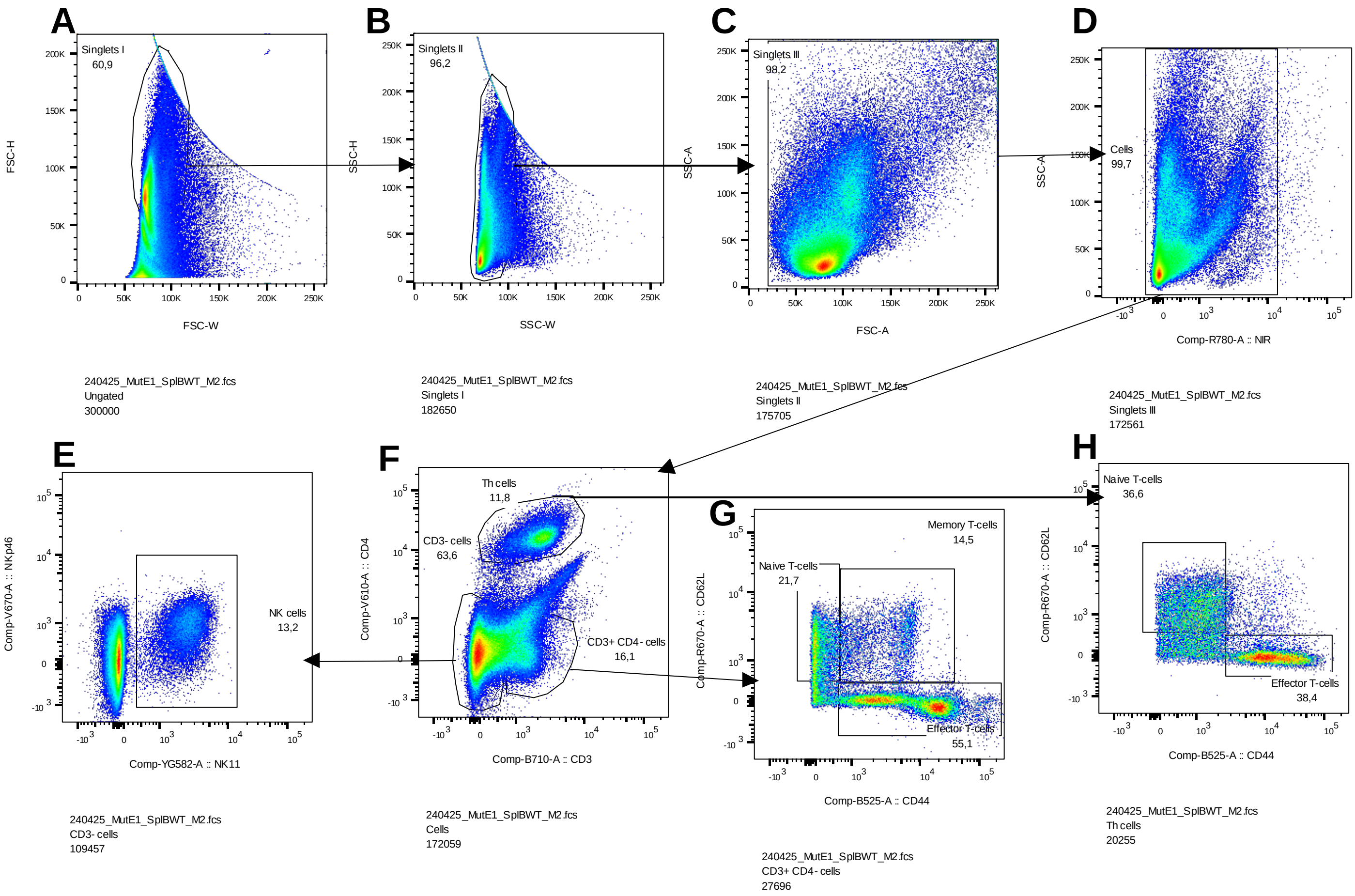

### Supplement Figure E5

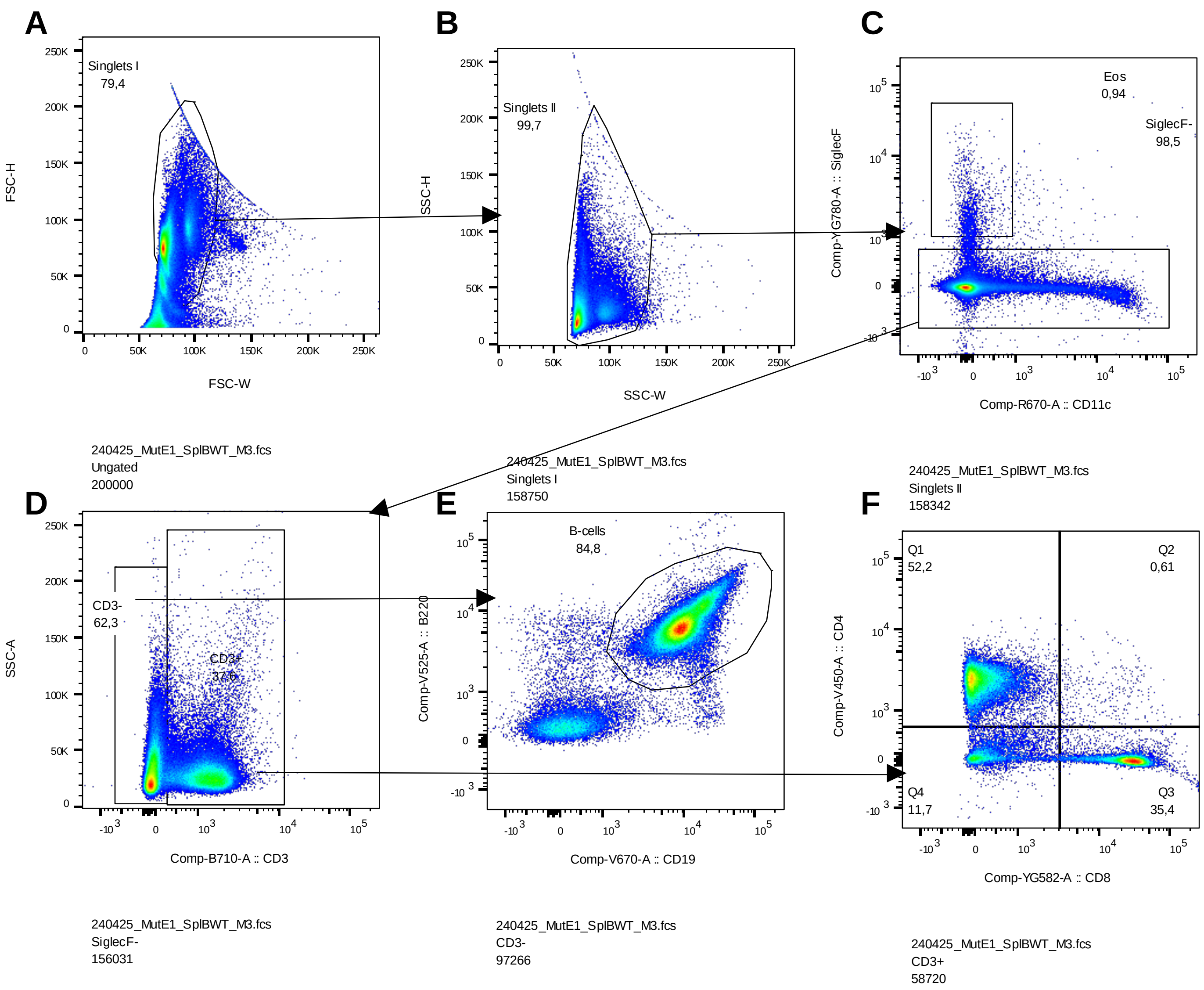

### Supplement Figure E6

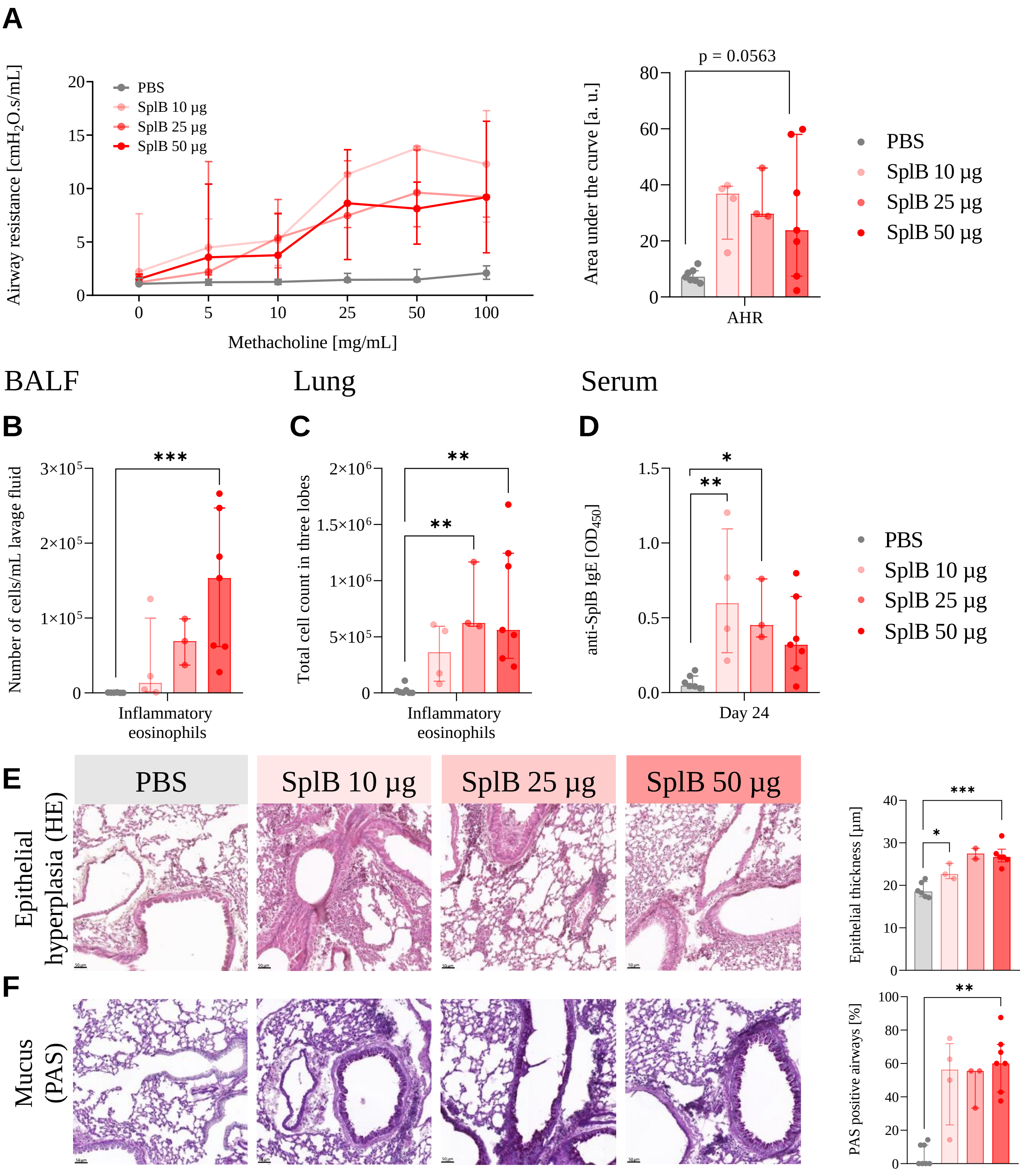

### Supplement Figure E7

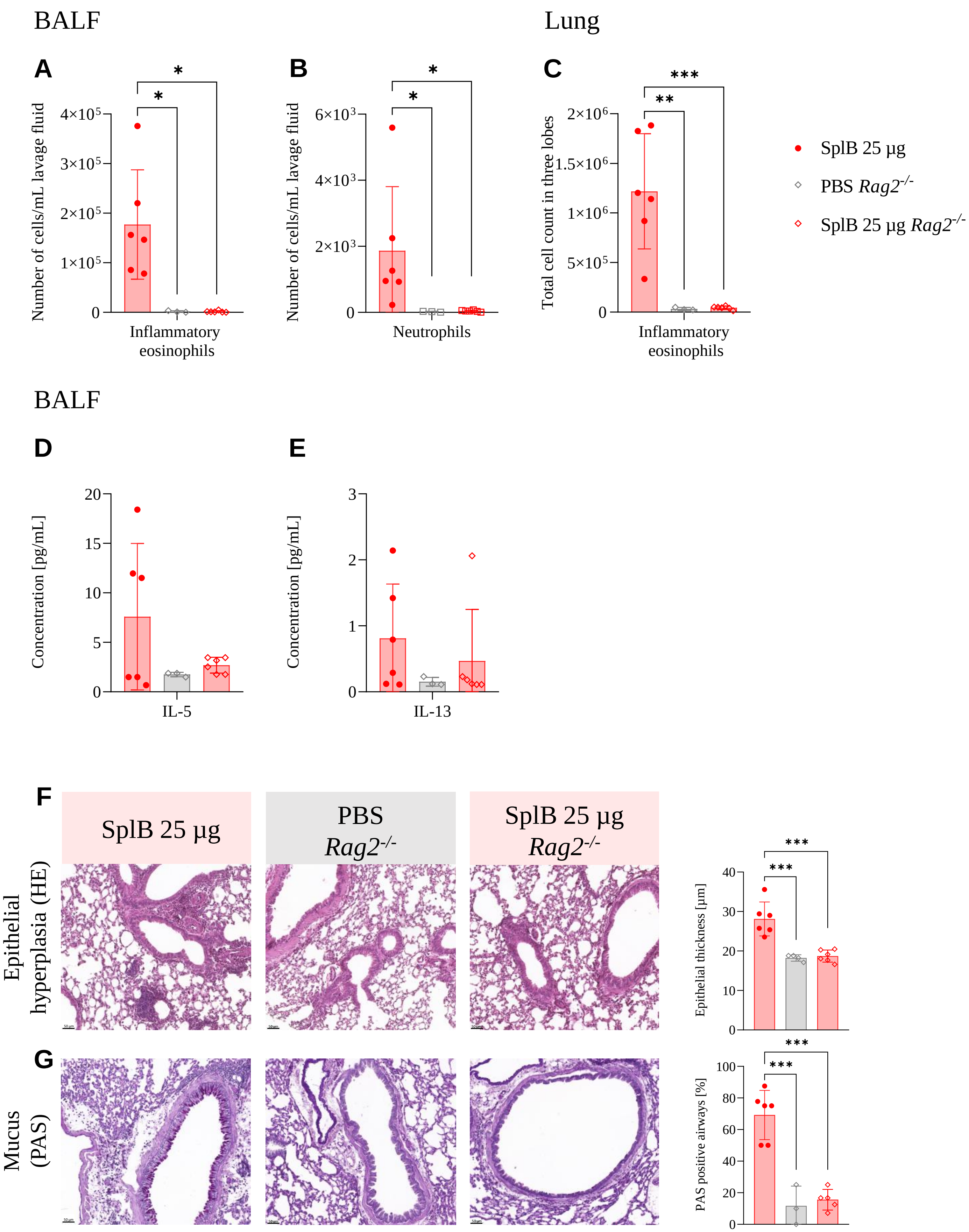

### Supplement Figure E9

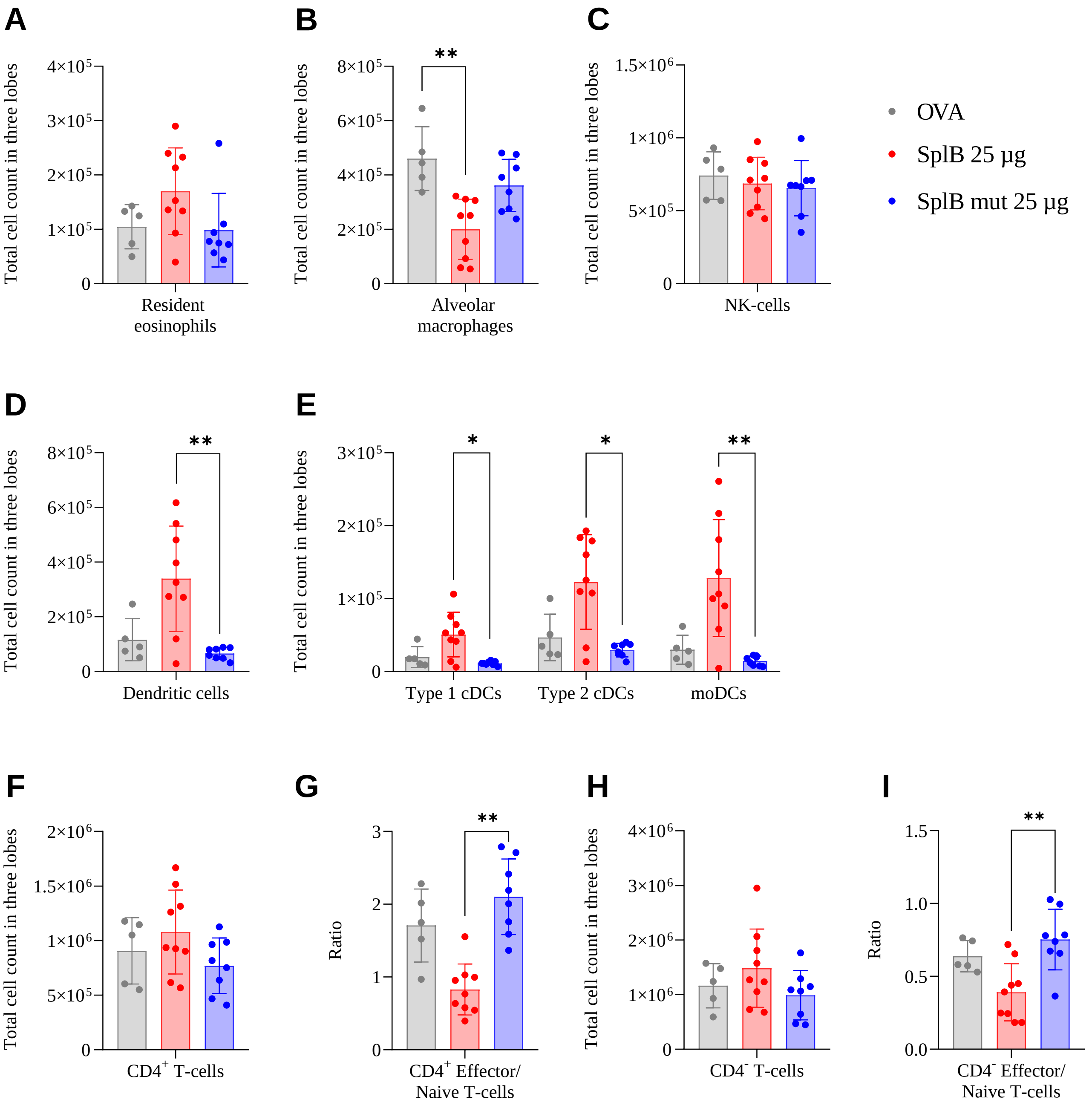

### Supplement Figure E10

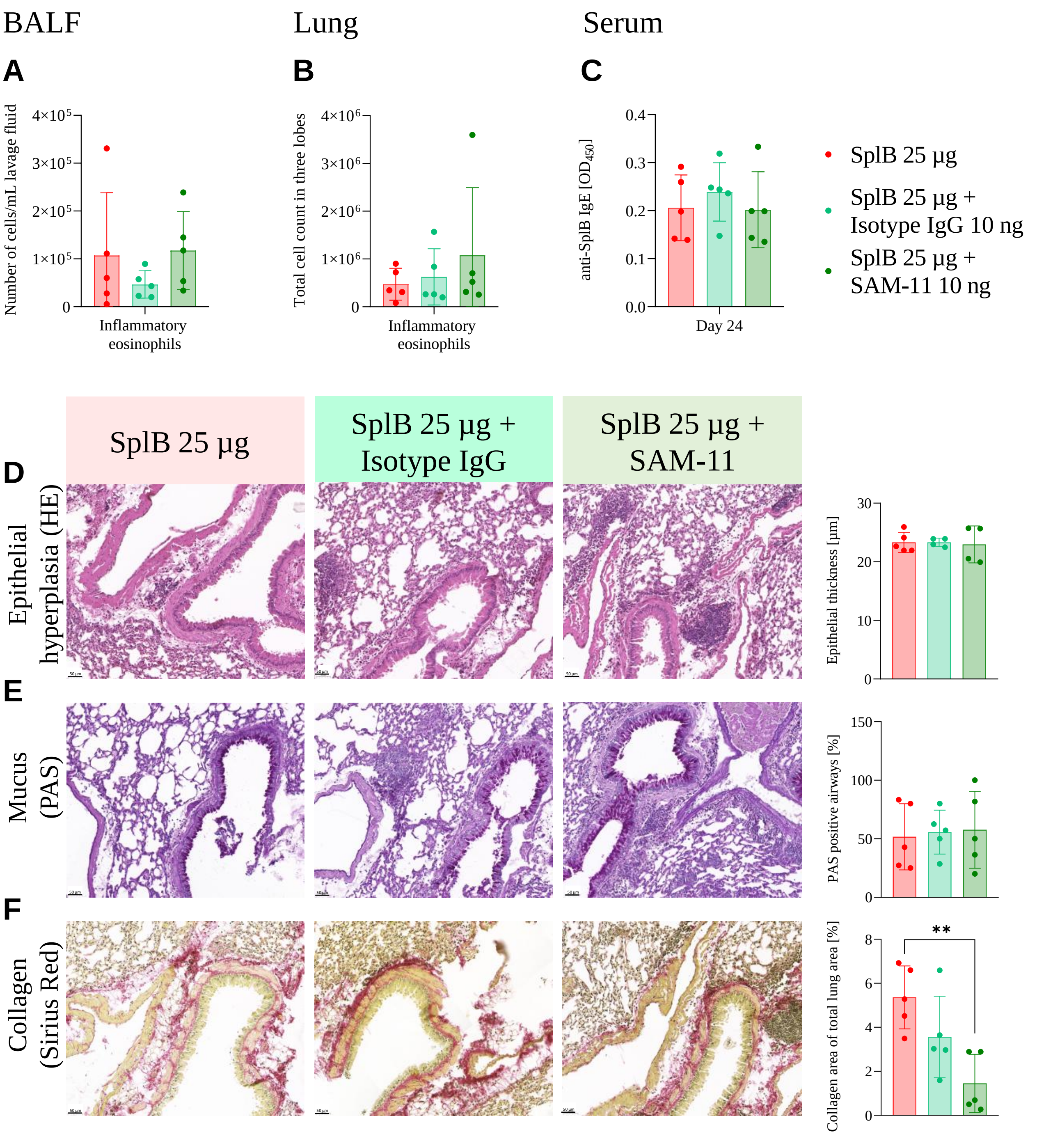
