## Supplement Figure E8 for "The *Staphylococcus aureus* serine protease-like protein B is a potent allergen in a murine asthma model"

**A**

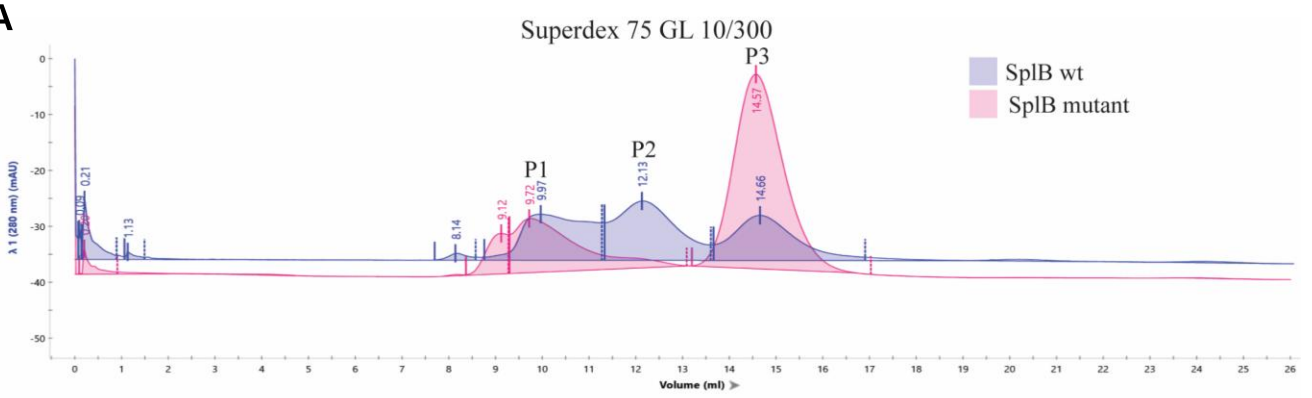

**B**

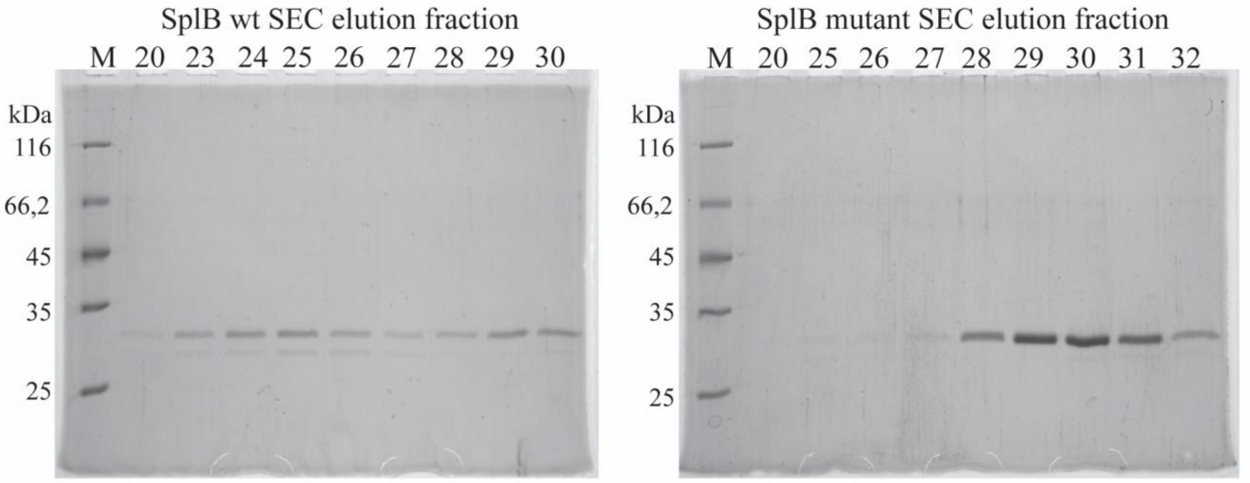

**C**

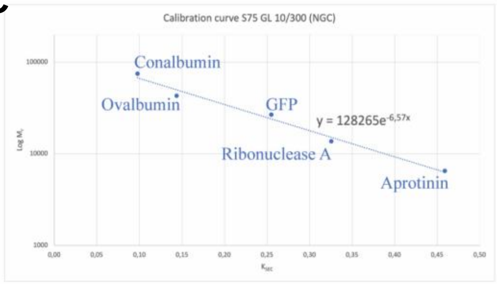

| | elution volume (mL) | apparent $M_r$ (Da) | apparent $M_r$ / theoretical $M_r$ |
|---|---|---|---|
| SplB_wt_peak 1 | 9.97 | 93464 | 3.58 |
| SplB_wt_peak 2 | 12.13 | 49334 | 1.89 |
| SplB_wt_peak 3 | 14.66 | 23341 | 0.89 |
| SplB_mut_peak 1 | 9.72 | 100638 | 3.86 |
| SplB_mut_peak 3 | 14.57 | 23971 | 0.92 |
