## Supplementary Methods for "The *Staphylococcus aureus* serine protease-like protein B is a potent allergen in a murine asthma model"

Online Repository

**Methods**

Recombinant proteins

Recombinant SplB was produced in the *S. aureus* strain RN4220 using the expression vector pTripleTREP^E1^. The sequence of SplB was derived from the *S. aureus* strain USA300 FPR3757 (Fig. E1 in the Online Repository). An amino acid switch (S193A) was introduced into the catalytic triad of SplB to generate a SplB mutant (SplB mut) without catalytic activity. Lipopolysaccharide contamination was monitored using the Endosafe Portable Test System (Charles River, USA). The quality of the preparations of active SplB and SplB mut was assessed by mass spectrometry analysis^E1^. Recombinant tag-free SplB was expressed in *B. subtilis* as described previously^E2^. Recombinant endotoxin-free ovalbumin (OVA) was purchased (InvivoGen, USA; vac-pova). Recombinant HpARI2 was kindly provided by Dr. Henry McSorley (Institute for Cell Signalling and Immunology, University of Dundee, UK).

Enzyme assays

A substrate-specific cleavage assay with a short peptide was performed to measure the catalytic activity of the SplB preparations. The synthetic peptide Ac-VEID-AMC (PeptaNova GmbH, Germany), which contains a validated SplB cleavage site^E3^, was used in this assay. 2.5 µM SplB was incubated with 25 µM peptide at 37 °C. Fluorescence intensity was quantified using a TECAN Infinite M200 plate reader (Tecan Group Ltd., Switzerland).

To test whether SplB is capable of cleaving murine IL-33 or the human complement factor C3 (hC3), enzymatically active SplB or SplB mut were incubated with murine IL-33 (OriGene Technologies, Inc., US; TP607659) or human C3 (Sino Biological Europe GmbH, Germany; 13182-H08H) (37 °C, 24 h). Equimolar concentrations of protease and substrate were used. Cleavage was monitored by Western blotting. 0.1 µg of the mIL-33 or hC3 digest were loaded on a 10% SDS gel and separated by SDS-PAGE. Afterwards, proteins were blotted onto a PVDF membrane. Goat anti-mouse IL-33 (R&D Systems, USA; AF3626) or rabbit anti-human C3 (ThermoFisher Scientific, USA; PA5-21349) was used in combination with HRP-conjugated donkey anti-goat IgG (R&D Systems, US; HAF109) or goat anti-rabbit IgG (OriGene Technologies, Inc., US; R1364HRP) for target detection. Bands were visualized with the enhanced chemiluminescence Maximum Sensitivity Substrate (ThermoFisher Scientific, USA).

Mice

Animals were maintained in a 12-hour/12-hour light/dark cycle and had access to water and food *ad libitum*. Animal experiments were approved by the responsible authority, the Landesamt für Landwirtschaft, Lebensmittelsicherheit und Fischerei Mecklenburg-Vorpommern (LALLF MV; AZ 7221.3-1-037/20). All animal experiments were carried out in accordance with the European Union Directive 2010/63/EU, the German Animal Welfare Act (Tierschutzgesetz), and the German Animal Welfare Ordinance on the Protection of Animals Used for Experimental Purposes (TierSchVersV). The following mouse strains were used: C57BL/6J, C57BL/6J-Rag2^em3Lutzy^/J (*Rag2^-/-^*), C57BL/6J *Il33^Gt/Gt^* (*Il33^-/-^*)^E4^ and B6.Cg-F2rl1tm1Mslb/J (*F2rl1^-/-^*)^E5^. Mice were purchased from Janvier (C57BL/6J), JAX (C57BL/6J *Rag2^-/-^* (strain 033526); C57BL/6J *F2rl1^-/-^* (strain 004993)) or bred in-house (C57BL/6J
*Il33^-/-^*, originally from Texas A&M Institute for Genomic Medicine (USA)). The mice were certified as *S. aureus*-free. All mice used in animal experiments were female and 8-12 weeks old when treatment started.

Mouse model of allergic airway inflammation (AAI)

Mice were put under anesthesia (75 mg/kg ketamine + 5 mg/kg xylazine) and treated intratracheally (i.t.) with PBS, SplB (10, 25 or 50 µg), SplB mut (25 µg) or OVA (50 µg) on days 0, 7, 14 and 21. Notably, no adjuvant or intraperitoneal (i.p.) priming was used in either of the groups. Proteins were dissolved in 50 µL PBS. In some experiments, the IL-33 inhibitor HpARI2 (10 µg), the PAR2-blocking mAb SAM-11 (10 ng; ThermoFisher Scientific, USA; 35-2300) or a monoclonal IgG2a isotype-antibody (10 ng; ThermoFisher Scientific, USA; 02-6200) were i.t. co-applied with SplB. On days 0, 7 and 14, approximately 100 µL blood was taken from the retroorbital plexus prior to the i.t. application. 72 hours after the fourth treatment, mice were anesthetized and the airway hyperreactivity (AHR) was measured. Afterwards the mice were sacrificed with an overdose of isoflurane.

Airway hyperreactivity test

AHR was measured by the forced oscillation technique using a FlexiVent system (Scireq, Canada). In deep anesthesia (75 mg/kg ketamine + 5 mg/kg xylazine) the animals’ trachea was exposed by surgical incision in the throat area. A small horizontal incision was made in the trachea and a canula was inserted. Afterwards, the mouse was connected to the FlexiVent and mechanically ventilated (3 cm H_2_O, 150 breaths per minute). Spontaneous breathing was prevented by i.p. injection of 2,5 mg/kg bodyweight rocuronium bromide (Inresa Arzneimittel GmbH, Germany). Acetyl-β-methyl-choline (methacholine) (Sigma-Aldrich, Germany) was aerosolized and delivered at increasing doses (0, 5, 10, 20, 50, 100 mg/mL) through inhalation. Airway resistance was measured throughout the next two minutes after each application.

Organ processing

Blood was collected, centrifuged (16,000× g, 4 °C, 10 min), and the serum was stored at -80 °C. A broncho-alveolar lavage was performed by injecting 1 mL of ice-cold PBS into the airways and slowly retracting it with a syringe. The procedure was repeated three times, the broncho-alveolar lavage fluid (BALF) was centrifuged (300 × g, 10 min, 4 °C), and the cell-free supernatant was taken off and stored at -80 °C. Cells were resuspended in 500 µL PBS and stained for flow cytometry analysis. Lungs were perfused with 5 mL PBS through the right ventricle. The upper part of the left lung lobe was fixed in 4% formalin for histology. The lower part of the left lung lobe and the postcaval lobe were snap-frozen in liquid nitrogen. The right superior, middle and inferior lobes were kept in RPMI at 4 °C. These lobes were digested with 80 μg/mL liberase (Sigma-Aldrich, Germany; 05401020001) and 320 μg/mL of DNase (Sigma-Aldrich, Germany; 10104159001) in 2.5 mL RPMI medium. Lobes were minced using the GentleMACS system (Miltenyi Biotec, Germany). The resulting cell suspension was then filtered through a 70 µm cell strainer to remove remaining fibers. After erythrocyte lysis, cells were resuspended in RPMI medium and stained for flow cytometry. Spleens were meshed through a 70 µm cell strainer to obtain single cell suspensions. Cells were washed with RPMI medium and centrifuged (300× g, 10 min, 4 °C). After erythrocyte lysis, cells were resuspended in RPMI medium for flow cytometry. Lung tissue from postcaval lobe was homogenized in RIPA lysis buffer supplemented with protease inhibitors using zirconium oxide beads (Bertin Technologies, France) in a Precellys 24 (6000 rpm; 2x20 s) (Bertin Technologies, France). The supernatant was collected. Protein concentration was determined by BCA assay (ThermoFisher Scientific, USA; 23225).

Flow cytometry

First, the cells from BALF, lung and spleen were counted on the BD LSR II flow cytometer (BD, USA) using TruCount beads (BD, USA). For phenotyping 1 million cells were incubated with Fc block solution (Miltenyi Biotech, Germany) and stained with a fixable live/dead stain (BioLegend^®^, USA). Antibody panels and gating strategies are provided in the supplement (Figs. E2-E5 in the Online Repository). Unless stated otherwise, all antibodies were purchased from Miltenyi (Miltenyi Biotech, Germany). Data were acquired using a BD LSR II flow cytometer and analyzed with the FlowJo software (V10.10.0). Spleen eosinophils are presented as percentages of total living cells instead of absolute numbers because only a portion of the spleen tissue was available for analysis, which precluded reliable calculation of absolute cell numbers.

Histology

Paraffin-embedded lungs were cut into 4 µm sections and stained with hematoxylin and eosin (HE), periodic acid-Schiff (PAS) (Carl Roth, Germany; HP01.1) or Sirius Red (Morphisto, Germany; 13425) according to the manufacturers’ instructions. The Congo red staining was performed by firstly immersing the deparaffinized sections in 0.5% alcoholic Congo red solution (0.5 g Congo red (Sigma-Aldrich, Germany; C6767-25G) in 100 mL 50% ethanol) for 30 minutes. Subsequently, the sections were differentiated in alkaline ethanol for a few seconds and immersed in Mayer's hematoxylin solution (Carl Roth, Germany; T865.1) to counterstain for 5 minutes. Sections were then dipped a few times in 0.05% lithium carbonate solution until sections turned blue, and washed in running tap water. For evaluation of eosinophil influx, eosinophils, stained in red, were counted manually in three different fields of view per lung section. One field of view was defined as 0.4 mm^2^. Slides were scanned with a Pannoramic MIDI II (Sysmex, Germany) and analyzed using 3DHistech CaseViewer Version 2.3 (3DHISTECH Ltd., Hungary). Sections of HE, PAS and Sirius Red staining are shown at 20× magnification, sections stained with Kongo Red are shown in 40× magnification.

Measurements of antibodies, albumin and cytokines

SplB- and OVA-specific IgE and IgG were measured with ELISA as described previously^E2^. In brief, 96-well plates were coated with 2 µg/mL tag-free SplB or OVA. After ON incubation at 4 °C, plates were blocked using 10% FCS in PBS (1 h, RT). For the IgE ELISA, the mouse serum was diluted 1:10, whereas for the IgG ELISA it was diluted 1:10,000. 50 µL of the serum samples were applied to each well and incubated (1 h, RT). After washing with PBS/Tween20 (0.05%), 50 µL of either rat-anti-mouse IgE (ThermoFisher Scientific, USA; MA5-16779), rat anti-mouse IgG (Jackson ImmunoResearch, USA; 115-065-164), rat anti-mouse IgG1 (BioLegend^®^, USA; 406603), rat anti-mouse IgG2a (BioLegend^®^, USA; 407103), rat anti-mouse IgG2b (BioLegend^®^, USA; 406703) or rat anti-mouse IgG3 (BioLegend^®^, USA; 406803) were used at a concentration of 1 µg/mL in 10% FCS in PBS for the detection step (1 h, RT). Following washing in PBS/Tween20 (0.05%), the plate was incubated with 50 µL of 0,33 µg/mL horseradish peroxidase (HRP) (Dianova, Germany; 016-030-084) (1 h, RT), after which it was washed again. TMB substrate (BD Biosciences, Germany) was added and the reaction was stopped after 20 min of incubation using 2N H_2_SO_4_. Absorbance was measured at 450 nm on a TECAN Infinite M200 plate reader (Tecan Group Ltd., Switzerland).

The total IgE (BioLegend^®^, USA; 432401) concentration in the serum and serum albumin (Antibodies.com, UK; A4028) concentration in the BALF were measured by ELISA according to the manufacturers’ instructions. The concentrations of cytokines (IL-33, TSLP, GM-CSF, IL-1α, IL-25) were quantified using kits from ThermoFisher Scientific (USA; IL-33, 88-7333-88) and BioLegend^®^ (USA; TSLP, 434104; GM-CSF, 432204; IL‑1α, 433401; IL-25, 447104) according to the manufacturers’ instructions. Cytokine concentration was measured in 50 µL mouse lung homogenate containing 50 µg of total protein.

Two bead-based cytokine detection kits were used to quantify cytokines and chemokines in the BALF (BioLegend^®^, USA; 741043 and 740007). Analysis was performed using the Qognit website (version: 2024-06-15).

In vitro epithelial barrier assays

Calu-3 human airway epithelial cells were cultured in MEM supplemented with 10% fetal calf serum (FCS), 1% penicillin/streptomycin, and L-glutamine. Cells were maintained under standard culture conditions at 37 °C and 5% CO₂.

For barrier integrity experiments, Calu-3 cells were seeded in triplicates at a density of 0.5 × 10⁶ cells per insert in 200 µL medium in Transwell inserts (costar, USA; 3470), with 800 µL medium added to the basolateral compartment. Medium was exchanged every 2–3 days. On day 10 after seeding, transepithelial electrical resistance (TEER) was measured to confirm epithelial confluence. Cells were stimulated apically with either 20 µg/mL SplB or the enzymatically inactive SplB mutant respectively, or medium alone as control. Stimulation was performed for 24 h at 37 °C. After stimulation, TEER was measured again to assess changes in barrier integrity. The experiment was repeated three times under identical conditions.

For assessment of epithelial cytotoxicity, Calu-3 cells were seeded in triplicates in flat-bottom 96-well plates at a density of 0.5 × 10⁶ cells per well in 200 µL complete medium. Medium was exchanged every 2–3 days. On day 10, cells were stimulated with SplB (20 µg/mL), enzymatically inactive SplB mutant (20 µg/mL), or medium alone as control. After 24 h incubation at 37 °C, LDH release was quantified using a commercial LDH cytotoxicity detection kit according to the manufacturer’s instructions (Sigma-Aldrich, Germany; 04744934001). The experiment was repeated three times under identical conditions.

Ex vivo human basophil stimulation

Human basophils were isolated from EDTA-anticoagulated peripheral blood obtained from healthy volunteers after written informed consent and approval by the local Ethics Committee (approval no. BB014/14). Blood samples were processed by density gradient centrifugation using Pancoll (PAN-Biotech GmbH, Germany; P04-601000). In brief, peripheral blood mononuclear cells (PBMCs) were isolated by carefully layering diluted full blood over Pancoll in 50 mL conical tubes. Samples were centrifuged at 800 × g for 30 min without brake. Following centrifugation, the PBMC layer at the interface was carefully collected. Cells were washed three times with DPBS by sequential centrifugation steps (200× g, 10 min, RT). Subsequently, basophils were purified using the Human Basophil Isolation Kit II (Miltenyi Biotec, Germany; 130-092-662) according to the manufacturer’s instructions. Cell purity was assessed by flow cytometry and consistently exceeded 80%. For purity assessment, cells were stained with anti-human CD303-FITC (Miltenyi Biotec, Germany, 130-113-192) and CD123-PE (Miltenyi Biotec, Germany, 130-113-326).

For functional assays, purified basophils were cultured in medium containing 5% FCS and 10 ng/mL IL-3 to support basal priming. Cells (1 × 10⁵ per condition in 25 µL) were seeded into 96-well plates and stimulated by adding 25 µL of stimulus solution. Basophils were incubated at 37 °C for 24 h with either 20 µg/mL SplB or an enzymatically inactive SplB mutant. Medium alone served as a negative control. Following stimulation, cells were harvested and centrifuged (300 × g, 10 min). Supernatants were collected for cytokine analysis using CBA (BioLegend^®^, USA; 741028). Cell pellets were used for flow cytometric analysis of activation markers. The following antibodies were used: anti-human CD203c-PE (BD Biosciences, Germany; 562972), CD63-APC (BioLegend^®^, USA; 353008), and FcεRI-PE/Cy7 (BioLegend^®^, USA; 334620).

Analytical size exclusion chromatography (aSEC)

aSEC runs were performed using a Superdex 75 GL 10/300 column on an NGC Discover FPLC system (Bio-Rad Laboratories; USA). 120 ml of a 100 mM SplB protein solution was injected onto the column and run through the column with a constant flow rate of 0.3 ml/min. Protein populations were separated during the chromatography and elution fractions of 500 ml were collected up to a total volume of 26 ml. Elution fractions containing protein were analysed by SDS-PAGE. The molecular weight (Mr) and oligomeric state of proteins in peak fractions were calculated using a calibration curve. The standard proteins Aprotinin, Ribonuclease A, GFP, Ovalbumin and Conalbumin were used to determine the elution volumes (Ve) of known molecular weights. The void volume (Vo) of the column was determined by dextran blue and the internal protein accessible volume (Vi) of the column was determined by b-mercaptoethanol. By using the equation KSEC = (Ve-Vo)/(Vi-Vo), the distribution coefficient (KSES) of the column was determined. KSEC values were plotted as a function of logMr and fitted using a logarithmic equation. The logarithmic equation was used to determine the apparent molecular weight of SplB. The calculation of the ration of apparent molecular weight and theoretical molecular weight gives an estimation of the oligomeric state of the protein.

Statistics

All datasets were tested for normality using the Shapiro–Wilk test. Ordinary one-way ANOVA followed by Tukey’s multiple comparisons test was used if all datasets were normally distributed, otherwise the Kruskal–Wallis test followed by Dunn’s multiple comparisons test was applied. AHR measurements were analyzed by calculating the area under the curve (AUC) and compare the AUC using the Kruskal–Wallis test followed by Dunn’s multiple comparisons test. A p value < 0.05 was considered significant. Statistical analyses were performed with GraphPad Prism v8.0.1. Data are shown as median with interquartile range if more than one group consisted of less than 4 animals, and as mean ± SD otherwise.
